# A Novel PHOX/CD38/MCOLN1/TFEB Axis Important For Macrophage Activation During Bacterial Phagocytosis

**DOI:** 10.1101/669325

**Authors:** Mehran Najibi, Joseph A. Moreau, Havisha H. Honwad, Javier E. Irazoqui

## Abstract

Macrophages are a key and heterogenous class of phagocytic cells of the innate immune system, which act as sentinels in peripheral tissues and are mobilized during infection. Macrophage activation in the presence of bacterial cells and molecules entails specific and complex programs of gene expression. How such triggers elicit the gene expression programs is incompletely understood. We previously discovered that transcription factor TFEB is a key contributor to macrophage activation during bacterial phagocytosis. However, the mechanism linking phagocytosis of bacterial cells to TFEB activation remained unknown. In this article, we describe a previously unknown pathway that links phagocytosis with the activation of TFEB and related transcription factor TFE3 in macrophages. We find that phagocytosis of bacterial cells causes an NADPH oxidase (PHOX)-dependent oxidative burst, which activates enzyme CD38 and generates NAADP in the maturing phagosome. Phago-lysosome fusion brings Ca^2+^ channel TRPML1/MCOLN1 in contact with NAADP, causing Ca^2+^ efflux from the lysosome, calcineurin activation, and TFEB nuclear import. This drives TFEB-dependent expression of important pro-inflammatory cytokines, such as IL-1α, IL-1β, and IL-6. Thus, our findings reveal that TFEB activation is a key regulatory event for the activation of macrophages. These findings have important implications for infections, cancer, obesity, and atherosclerosis.

## Introduction

Macrophages are phagocytic cells of the innate immune system, which act as sentinels in peripheral tissues and are mobilized during infection ^1^. Macrophages are activated in the presence of bacteria, which entails specific programs of gene expression. Depending on the stimulus and the microenvironment, particularly the pathogen molecules and cytokines present, macrophages adopt phenotypes (or “polarize”) along a spectrum, from “classical” activation (a.k.a. M1 polarization), which is predominantly pro-inflammatory, to “alternative” activation (a.k.a. M2 polarization), which is predominantly pro-resolution ^2^. The activation state of the macrophage has great functional consequences for health and disease.

Because transcriptional regulation contributes greatly to the macrophage phenotype, understanding the transcription factors that are involved is of paramount importance. Similarly, it is imperative to fully elucidate the signaling pathways that regulate such transcription factors under conditions of homeostasis and disease. Much work has characterized important signaling from the Toll-like receptors (TLRs), which recognize molecules produced by pathogens such as bacterial cell wall components and trigger the activation of transcription factors NF-κB, IRF3, and AP1 ^3^. All three examples of transcription factors are subject to multiple regulatory layers, nuclear-cytoplasmic shuttling being a major mechanism of regulation.

We recently identified TFEB as a transcription factor that is important for cytokine and chemokine gene induction in macrophages following bacterial infection^4^. We found that TFEB resides in the cytosol in resting murine macrophages, but is imported to the nucleus after short infection with *Staphylococcus aureus* or *Salmonella* Typhimurium ^4,5^. Moreover, cells defective in TFEB expression exhibit defective induction of important pro-inflammatory cytokines, including TNF-α, IL-1β, and IL-6 ^4^. Subsequent studies confirmed our initial observations, and extended them by showing that TFE3 is also activated by lipopolysaccharide (LPS), a cell wall constituent of Gram-negative bacteria ^6^. Moreover, an independent study demonstrated how phagocytosis of opsonized particles triggers TFEB nuclear import and the subsequent activation of bactericidal mechanisms in murine macrophages ^7^. Thus, clearly activation of TFEB is an important regulatory event in macrophage responses to bacterial pathogens. However, the regulatory mechanisms upstream of TFEB during bacterial infection in macrophages remained unclear.

We recently reported a novel pathway that is required for TFEB activation during bacterial infection in macrophages ^5^. In it, phosphatidyl choline-directed phospholipase C (PC-PLC), which produces diacyl glycerol (DAG), activates protein kinase D1 (PrKD1) to promote TFEB nuclear import. While in the absence of this pathway TFEB cannot be activated by infection, its induction is sufficient to cause TFEB nuclear import. However, the molecular mechanism linking PrKD1 to TFEB remained unknown. Importantly, the link between phagocytosis of bacteria and PC-PLC/PrKD1 activation remained unclear.

In the present article, we describe the discovery of a novel pathway that links phagocytosis of bacterial cells to TFEB activation and downstream cytokine expression. Instead of the expected TLR signaling pathways, we found that signal transduction for TFEB activation requires NADPH oxidase (PHOX) and the production of reactive oxygen species (ROS). This event sets in motion the activation of CD38, which produces NAADP, an activating ligand for lysosomal Ca^2+^ export channel TRPML1/MCOLN1. TRPML1/MCOLN1 activation, followed by Ca^2+^ export, drives calcineurin, a Ca^2+^-regulated protein phosphatase that dephosphorylates TFEB on conserved S/T residues critical for TFEB cytosolic sequestration ^8^, thus allowing TFEB nuclear import and downstream gene induction.

Because of the broad tissue expression of CD38 and other components of this pathway ^9^, and because several stresses and diseases cause potentially TFEB-activating ROS production from mitochondria ^10–12^, the findings reported here are likely to have broad implications for the regulation of gene expression down-stream of TFEB in multiple tissues and physiological conditions, including homeostasis and inflammatory diseases.

## Results

### Phagocytosis of Gram+ and Gram-bacteria activates TFE3 in macrophages

We previously showed that *S. aureus* and *S.* Typhimurium could activate TFEB in murine macrophages ^5^. Since then, several studies have suggested that TFE3 and TFEB share many features of upstream regulation ^13–15^. In addition, previous work showed that extended incubation with LPS activates TFE3 in macrophages ^6^. Therefore, we wondered if TFE3 might also respond to infection by bacteria. We found that infection with *S. aureus* or *S.* Typhimurium activated TFE3 with similar kinetics and amplitude as TFEB, measured by its nuclear accumulation (**Fig. 1A-H** and **S1A-C, F**). Moreover, peptidoglycan (PGN) from *S. aureus* alone was sufficient to induce TFE3 and TFEB nuclear accumulation (**Fig. 1C, F-H**), suggesting that phagocytosis of large particles was not a requirement. Both *S. aureus* and PGN also induced the formation of lysosomes, measured with Lysotracker staining (**Fig. 1I-R**). There was a striking correlation between nuclear accumulation of TFEB and lysosome induction, indicating that *S. aureus-* or PGN-induced TFEB activation was biologically relevant.

**Figure 1.**
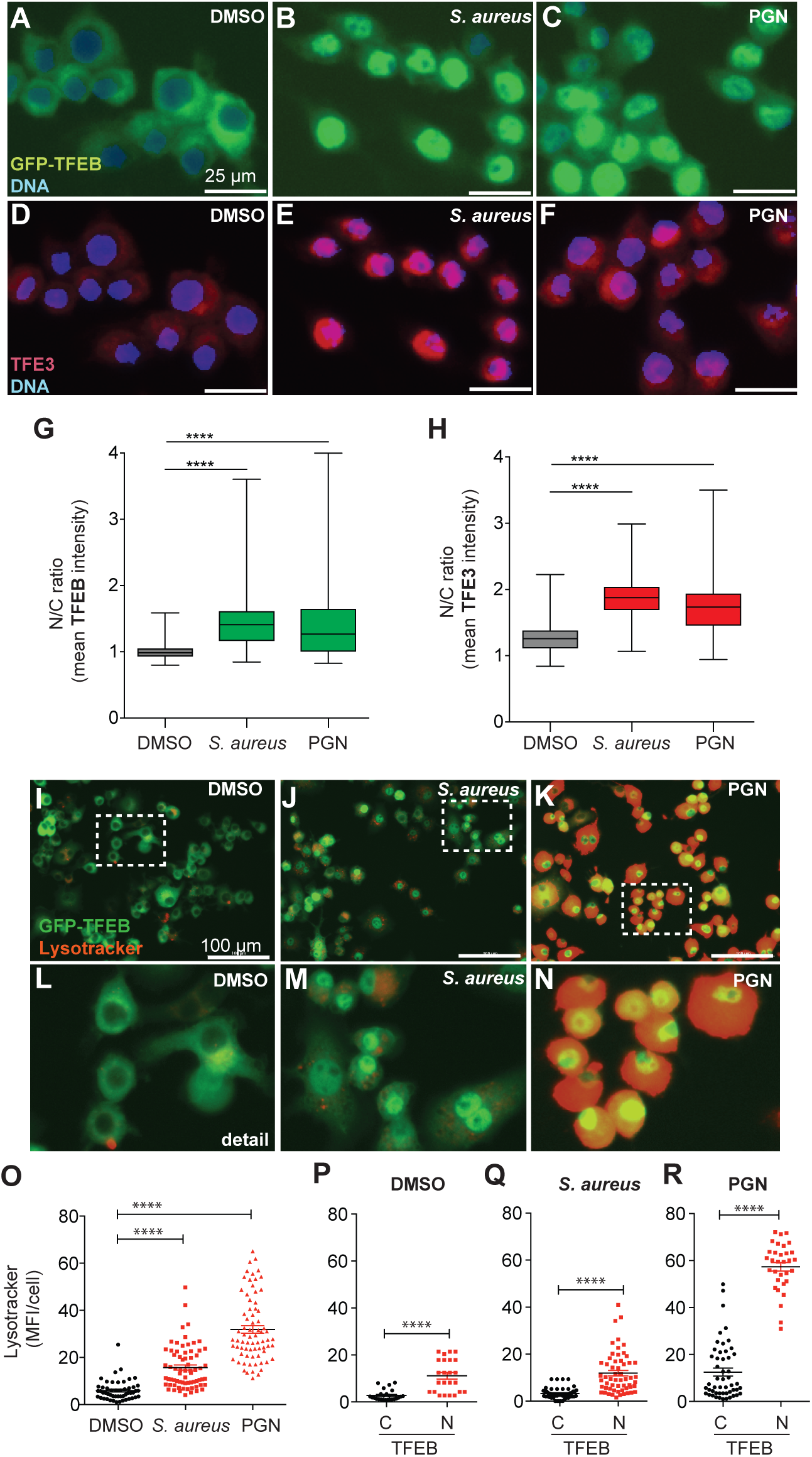
Gram+ bacterial cells and ligands activate TFE3. **A-H.** GFP-TFEB immortalized BMDM (iBMDMs) were infected with *S. aureus* (MOI = 10) for 3 h. Incubation with 10 µg/ml PGN from *S. aureus* was 3 h. Cells were fixed and processed for anti-TFE3 immunofluorescence staining. Shown are representative images, and quantification of three biological replicates (n = 450). **A**. DMSO control, **B**. *S. aureus* infection, and **C**. *S. aureus* PGN (10 µg/ml). **D**. DMSO control, **E.** *S. aureus* infection, **F**. *S. aureus* PGN. **G.** GFP-TFEB intensity in the nucleus compared to the cytoplasm (N/C ratio) measured using CellProfiler. **** p ≤ 0.0001 (one-way ANOVA followed by Tukey’s post-hoc test). (**H**) Anti-TFE3 fluorescence N/C ratio measured using CellProfiler. **** p ≤ 0.0001 (one-way ANOVA followed by Tukey’s post-hoc test). **I-N**. GFP-TFEB iBMDMs were infected with *S. aureus* or incubated with *S. aureus* PGN as above, and stained with Lysotracker. Shown are representative images and quantification of three biological replicates (n = 60). **I**. DMSO control. **J**. *S. aureus* infection, and **K**. *S. aureus* PGN. **L-N**. Higher magnification micrographs for DMSO control (**L**), *S. aureus* (**M**), and PGN (**N**). **O**. Lysotracker intensity per cell (MFI/cell) measured with CellProfiler. **** p ≤ 0.0001 (one-way ANOVA followed by Tukey’s post-hoc test). **P-R.** Comparison of Lysotracker intensities in cells with TFEB cytoplasmic localization (C) with that in cells with nuclear localization (N) following incubation with DMSO (**P**), *S. aureus* (**Q**), or PGN (**R**). **** p ≤ 0.0001 (two-sample two-tailed unpaired *t* test).

*Salmonella* infection produced a more nuanced response. As we previously showed, live and dead *Salmonella* differed in their ability to induce TFEB activation ^5^. While live *Salmonella* induced rapid activation of TFEB and TFE3, dead *Salmonella* did so much more slowly (**Fig. S1A-D, F, G**). Interestingly, the kinetics of TFEB and TFE3 activation by dead *Salmonella* resembled that of LPS, suggesting that they might trigger TFEB/3 activation through a similar sensing mechanism (**Fig. S1E, H**). In contrast to *S. aureus, Salmonella* did not induce the accumulation of lysosomes, but rather decreased it below baseline (**Fig. S2A**). LPS induced lysosomal accumulation, as PGN did (**Fig. S2C**). Thus, we suspect that *Salmonella* possesses a mechanism to inhibit lysosomal biogenesis down-stream of TFEB/3. This is consistent with previous results showing lysosomal inhibition by *Salmonella* ^16,17^.

The overall conclusion of these observations is that bacterial infection and bacterial ligand stimulation of murine macrophages causes biologically significant TFEB and TFE3 activation. However, the functional consequences of bacterially-induced TFEB and TFE3 nuclear import were unclear.

### TFEB and TFE3 are key contributors to the macrophage transcriptional response to infection

To better understand the functional consequences of TFEB and TFE3 activation during bacterial infection, we performed RNAseq of a RAW264.7 murine macrophage cell line harboring deletions in *Tfeb* and *Tfe3.* We detected induction of 1,022 genes after 3 h of *S. aureus* infection in wild type macrophages (**Table S1**, **Fig. 2A**). Compared to wild type cells, *Tfeb*^-/-^ *Tfe3^-/-^* (double knockout, or dKO) cells exhibited a drastically altered transcription profile (**Fig. 2A, C**). Over-all, about two-thirds of *S. aureus-*induced genes in wild type macrophages were not induced in dKO cells (**Fig. 2D**), indicating that a large majority of the transcriptional response to *S. aureus* was TFEB/3-dependent. In addition, while wild type macrophages induced *Ccl5, Nos2, Ptgs2,* and *Tnf,* indicating a classically-activated state, dKO cells were completely defective in their induction (**Fig. 2B**). This revealed that macrophage polarization to the classically activated state, or M1, requires TFEB and/or TFE3.

**Figure 2.**
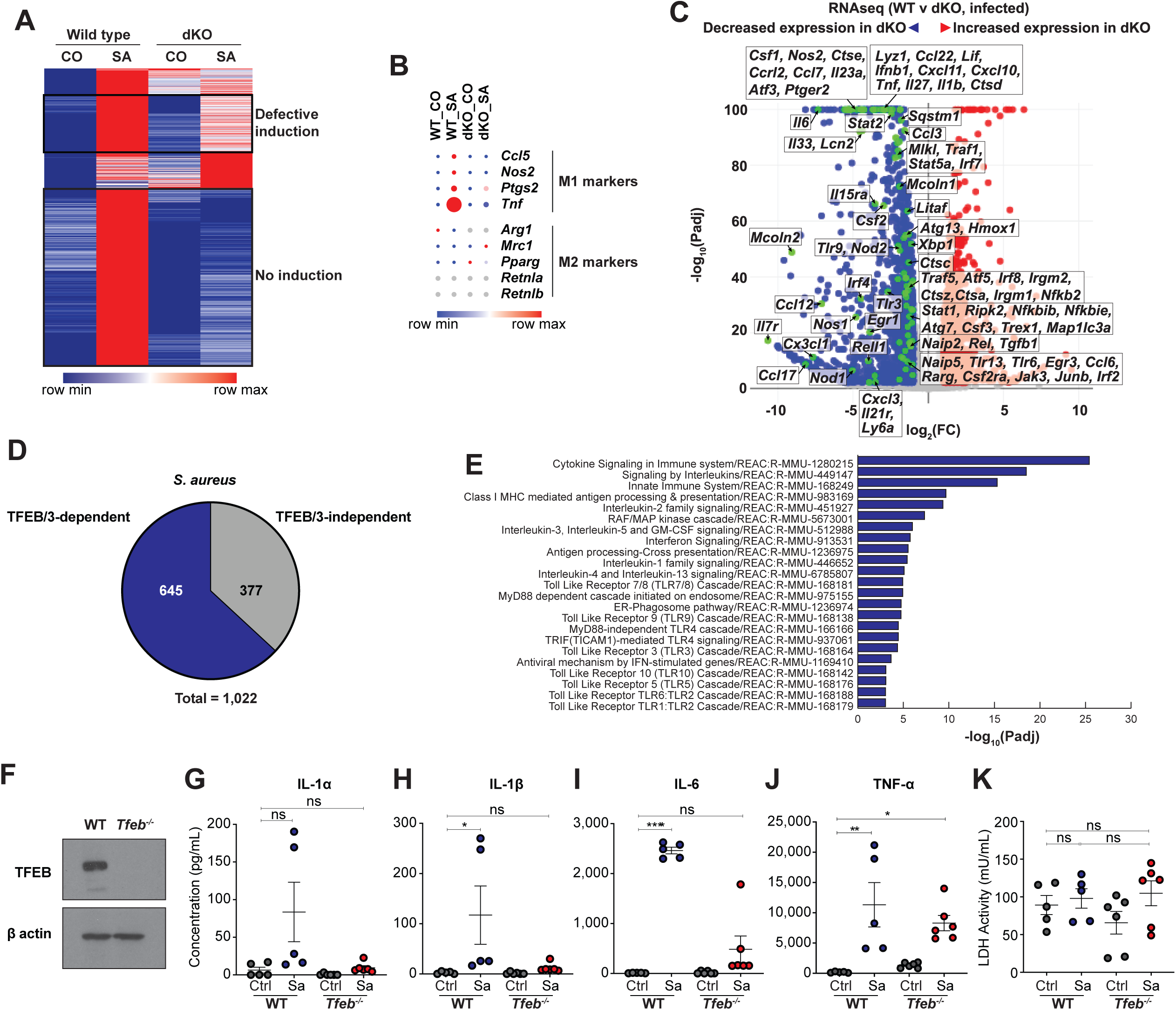
TFEB and TFE3 are required for the macrophage transcriptional response to *S. aureus.* **A-E**. Wild type (WT) and *Tfeb Tfe3* double knockout (dKO) RAW264.7 cells were infected as described in *Methods* for 3 h. N = 3 biological replicates. **A.** Heat map showing expression of genes that were differentially expressed in wild type cells. Shown are baseline (CO) and infected (SA) conditions, in both cell lines. Two groups of genes with defective induction in dKO cells are indicated with black boxes. **B.** Heat map showing expression of M1 and M2 marker genes. Colors indicate direction and magnitude of expression per row; circle diameter is proportional to absolute expression. Grey indicates no expression detected. **C.** Volcano plot showing differential gene expression in infected dKO cells relative to infected WT cells. Blue color denotes downregulation in dKO cells; red indicates upregulation in dKO cells. Green highlights noteworthy innate immunity genes, with corresponding adjacent labels. **D.** Pie chart showing percent of genes that are upregulated in infected v uninfected WT cells, and which are TFEB/3-dependent. **E.** Reactome pathway over-representation analysis of TFEB/3-dependent genes, plotting adjusted P values for each category (Padj). **F.** Verification of TFEB expression in *Tfeb*^flox/flox^ (WT) and *Tfeb*^ΔLysM^ (*Tfeb*^-/-^) iBMDMs by anti-TFEB immunoblot. **G-K**. Wild type (WT) and *Tfeb*^ΔLysM^ (*Tfeb*^-/-^) iBMDMs were incubated with PBS (Ctrl) or infected with *S. aureus* (Sa, MOI = 10). Concentrations of IL-1α (**G**), IL-1β (**H**), IL-6 (**I**), TNF-α (**J**), and lactate dehydrogenase activity (LDH, cytolysis control, **K**) were measured in culture supernatants 13 h following infection, by ELISA and LDH assay, respectively. N = 5-6, *p < 0.05, **p < 0.01, ***p < 0.001, ****p < 0.0001, one-way ANOVA followed by Tukey’s post-hoc test.

A large group of innate immunity genes exhibited decreased expression in dKO cells (**Fig. 2C**). Noteworthy examples included TLR genes *Tlr6, Tlr9,* and *Tlr13,* NF-κB transcription factor genes *Nfkb2* and *Rel,* NF-κB inhibitor genes *Nfkbib* and *Nfkbie,* and NLR genes *Nod1, Nod2, Naip2,* and *Naip5.* In addition, several cytokine genes exhibited defective expression, including *Tnf, Il1b, Il6, Ifnb1, Tgfb1, Csf1, Csf2, and Csf3,* and various interleukins (e.g. *Il23a, Il27,* and *Il33)*, as well as chemokines and chemokine receptors such as *Ccl3, Ccl6, Ccl7, Ccl12, Ccl17, Ccl22, Cxcl3, Cxcl10, Cxcl11, Ccrl2, Cx3cr1,* and signaling components such as *Ripk2, Jak3, Stat1, Stat2,* and *Stat5a.* Moreover, several autophagy and lysosomal genes showed lower expression: Cathepsins (lysosomal proteases *Ctsa, Ctsd, Ctse, Ctsz*), *Sqstm1, Mcoln1* and *Mcoln2, Atg7, Atg13, Irgm1* and *Irgm2,* and *Map1lc3a* (LC3). Finally, expression of ER unfolded protein response transcription factor gene *Xbp1* and necroptosis effector gene *Mlkl* were also reduced in dKO macrophages. These results strongly suggest that TFEB/3 are important for many key functions of macrophages, including pathogen recognition, pro-inflammatory signaling, secretion of chemotactic signals, and autophagy/lysosomal clearance.

More systematically, analysis of over-represented Reactome pathways ^18^ revealed that the top affected categories were “Cytokine Signaling in Immune System”, “Signaling by Interleukins”, and “Innate Immune System”. Other significant categories affected were “Interferon signaling”, “ER-Phagosome Pathway”, and various TLR pathways (**Fig. 2E**). Consistent with the transcriptome measurements, we observed robust secretion of IL-6 in *S. aureus-*infected wild type murine macrophages, which was abrogated by deletion of *Tfeb* alone (**Fig. 2F, H**). Wild type cells also secreted more IL-1β and TNF-α than *Tfeb*^ΔLysM^ macrophages, even though in this case the secreted levels were noisier than IL-6 (**Fig. 2H, J**). IL-1α exhibited a similar trend to IL-1β (**Fig. 2G**). Lactate dehydrogenase (LDH) release assays ruled out that these differences were due to cytotoxicity (**Fig. 2K**). Together, these observations provide strong evidence that TFEB/3 are essential for the macrophage transcriptional response to *S. aureus* infection, for macrophage polarization and function, and for the production of pro-inflammatory cytokines and chemokines. They also suggest that *Tfeb* deletion alone is sufficient to confer a defect in cytokine production, consistent with previous results ^6^. Therefore, nuclear translocation of TFEB and TFE3 is a key regulatory event for the macrophage response to infection. However, the signaling events upstream of TFEB/3 activation remained undefined.

### Bacterial infection activates TFEB and TFE3 independently of TLR signaling

Because TLRs constitute a major mechanism of innate recognition of bacterial ligands, we investigated the roles of TLR signaling in TFEB activation by bacterial cells and ligands. MyD88 is a protein adaptor that is important for signaling from TLR2/6 and TLR4, which detect PGN and LPS, respectively ^19^. TRIF is an adaptor that is also important for signaling from endosomal TLR4 ^20^.Together, MyD88 and TRIF are essential for TLR signaling. Therefore, we examined TFEB activation in murine macrophages lacking both MyD88 and TRIF.

Much to our surprise, after infection with *S. aureus* and treatment with PGN *Myd88^-/-^ Trif^-/-^* macrophages exhibited the same levels of TFEB activation as wild type cells (**Fig. 3**). Similarly, after infection with *Salmonella* or treatment with TLR4 agonists LPS or MPLA we observed no difference between *Myd88^-/-^ Trif^-/-^* and wild type macrophages (**Fig. S2**). These results suggested that TLR signaling is dispensable for TFEB activation by bacterial cells and ligands, regardless of cell wall structure.

**Figure 3.**
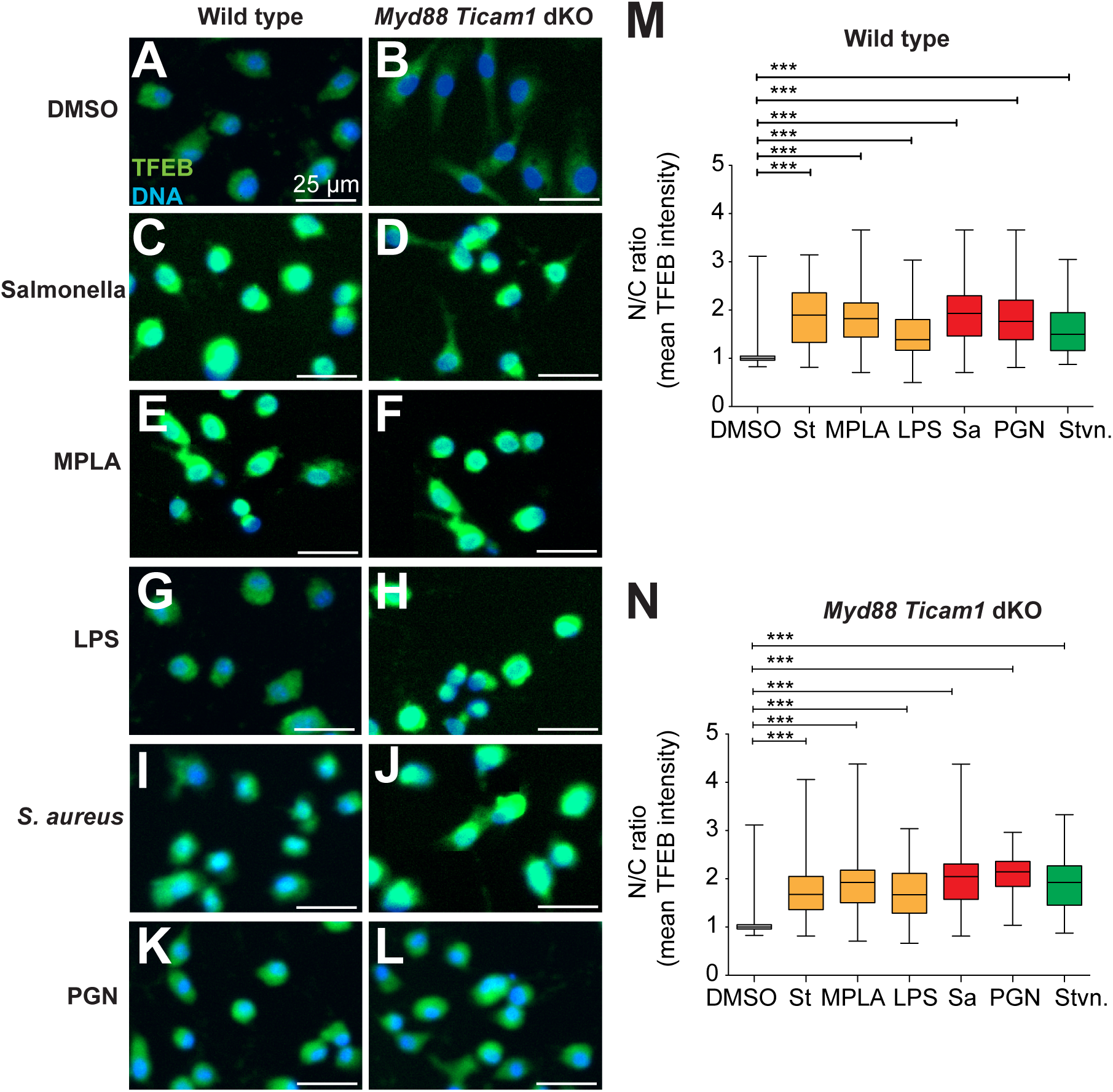
TLR signaling is dispensable for TFEB activation. **A-N**. Primary BMDMs from WT and *Myd88 Trif* double knockout mice (*Myd88 Trif* dKO) were infected, fixed, and processed for anti-TFEB immunofluorescence staining. TFEB fluorescence N/C ratio per cell was measured using CellProfiler (n = 250). Shown are representative images and quantification of three biological replicates. **A, B**. DMSO control (t = 6 h); **C, D.** *Salmonella* Typhimurium (St; MOI = 1, t = 6 h). **E, F.** Monophosphoryl lipid A (MPLA, 1 µg/ml, t = 6 h). **G, H.** *Salmonella* Typhimurium lipopolysaccharide (LPS, 1 µg/ml, t = 6 h). **I, J**. *S. aureus* (Sa, MOI = 10, t = 3 h). **K, L**. *S. aureus* peptidoglycan (PGN, 10 µg/ml, t = 3 h). **M, N.** Quantification of TFEB N/C ratio per cell in WT (**M**) and *Myd88 Trif* dKO (**N**) cells. *** p ≤ 0.001 (one-way ANOVA followed by Tukey’s post-hoc test).

To test this conclusion further, we examined TFEB activation in macrophages deficient for TLR2 and TLR4 (**Fig. S3A, B**). After infection with *S. aureus* and PGN or Pam3Csk4 treatment, *Tlr2*^-/-^ macrophages exhibited the same activation of TFEB as wild type cells (**Fig. S3C-L**). Likewise, after infection with live *Salmonella* or treatment with LPS, TFEB activation in *Tlr4*^-/-^ and wild type macrophages was indistinguishable; MPLA still activated TFEB in *Tlr4*^-/-^ cells, albeit more weakly (**Fig. S3M-X**). Together, these results indicated that bacterial cells and ligands activate TFEB by a TLR-independent pathway.

### Bacterial activation of TFE3 requires the PC-PLC/PrKD1 pathway

Given this lack of relevance of TLR signaling, we sought alternative mechanisms of TFE3 activation. We previously showed that TFEB activation in macrophages infected with *S. aureus* or *Salmonella* required the activity of a pathway upstream of PrKD1 ^5^. We found that prior inhibition of PC-PLC with D609 or of PrKD with kb-NB142-70 completely abrogated TFE3 activation by *Salmonella,* LPS, and *S. aureus,* similar to TFEB (**Fig. S4A-N**). Furthermore, we observed strong TFE3 activation by the DAG analog PMA, which was prevented by PrKD inhibition (**Fig. S4O, P**). Together, these data indicate that the PC-PLC/PrKD1 pathway is necessary and sufficient for TFE3 activation during infection in macrophages, as we previously showed for TFEB. However, the connection between PrKD1 and TFEB (or TFE3) nuclear import remained unclear. More importantly, the mechanism connecting phagocytosis of bacteria to TFEB/3 activation remained unknown. Therefore, we decided to examine regulatory events more proximal to TFEB.

### Bacterial activation of TFEB requires the TRPML1/MCOLN1 – Ca^2+^ – calcineurin pathway

The Ca^2+^-dependent protein phosphatase calcineurin was recently shown to be principally responsible for TFEB nuclear import induced by starvation in HeLa, HEK293, and human fibroblasts, as well as mouse muscle and MEFs. Such effect was mediated by its dephosphorylation of mTORC1 target sites in TFEB ^21^. However, the role of calcineurin in TFEB control in infected macrophages has not been tested before. Therefore, we evaluated TFEB activation in infected macrophages lacking calcineurin function. Prior incubation with Ca^2+^-chelating agent BAPTA prevented TFEB activation by *S. aureus* (**Fig. 4A-E**), suggesting that intracellular Ca^2+^ is important. Moreover, prior treatment with calcineurin inhibitor FK506 prevented *S. aureus-*triggered TFEB activation even more effectively (**Fig. 4D, E**), suggesting that calcineurin activity was required. Furthermore, siRNA-mediated silencing of genes *Ppp3cb,* which encodes calcineurin catalytic subunit β, or *Ppp3r1,* which encodes calcineurin regulatory subunit Bα, abrogated TFEB activation by *S. aureus* (**Fig. 4F-L**). These experiments indicate that calcineurin is required for TFEB activation during *S. aureus* phagocytosis. We found similar results during infection with *Salmonella* (**Fig. S5A-L**), indicating that calcineurin is a general requirement for TFEB activation during infection of macrophages. Furthermore, the Ca^2+^ ionophore ionomycin was sufficient to trigger TFEB nuclear import in uninfected macrophages (**Fig. 4M-O**). Thus, activation of calcineurin is necessary and sufficient for TFEB nuclear import in macrophages during infection with *S. aureus* or *Salmonella*.

**Figure 4.**
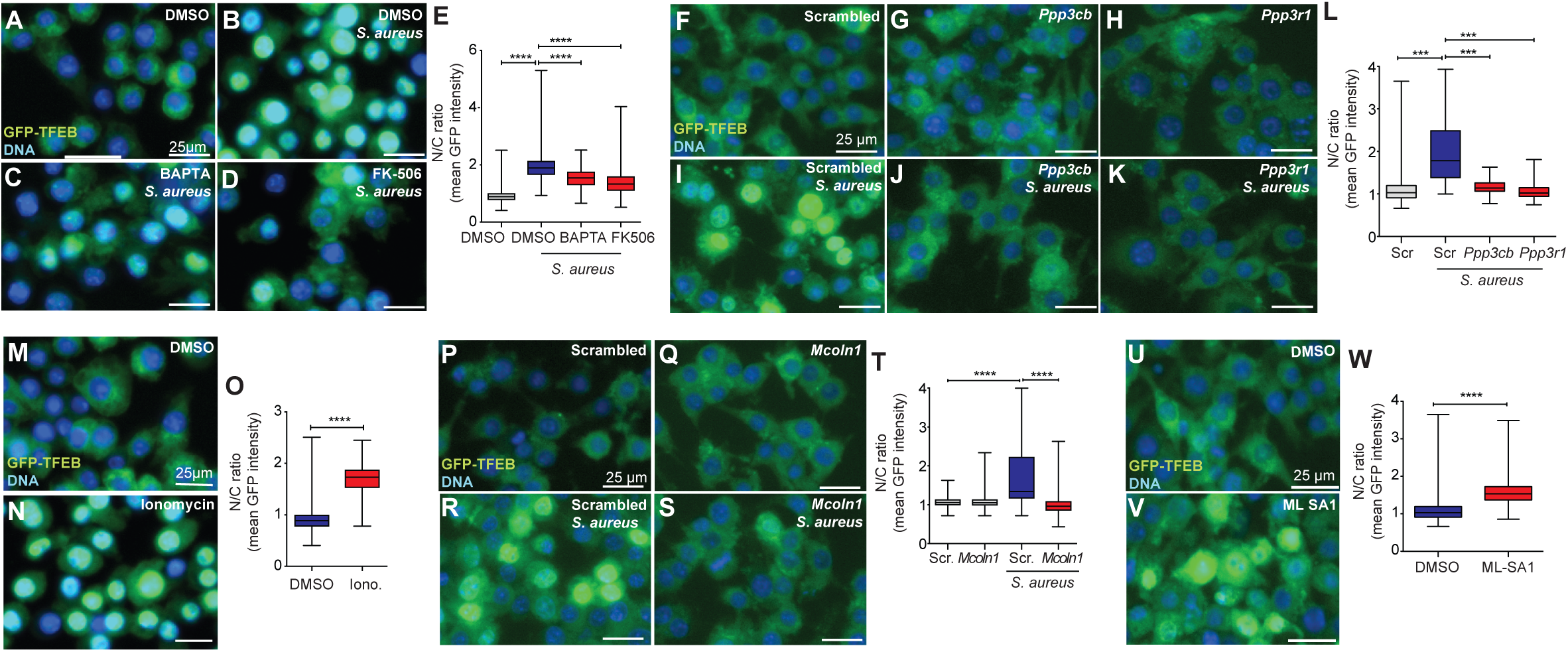
TFEB activation is mediated by TRPML1/MCOLN1, Ca^2+^, and calcineurin. **A-E.** GFP-TFEB iBMDMs were treated with DMSO without infection (t = 3 h, **A**) or treated for 3 h and subsequently infected with *S. aureus* (**B**, MOI = 10, t = 3 h). In parallel, cells were treated with BAPTA (**C,** 10 μM, t = 3 h) or FK506 (**D,** 5 μM, t = 6 h) and subsequently infected with *S. aureus* (MOI = 10, t = 3 h). **E.** Quantification of GFP-TFEB N/C Ratio by CellProfiler (3 biological replicates, n = 350 cells). **** p ≤ 0.0001 (one-way ANOVA followed by Tukey’s posthoc test). **F-K.** GFP-TFEB iBMDMs were treated with scrambled (Scr, **F**) or siRNA against *Ppp3cb* (**G**), *Ppp3r1* (**H**) for 48 h. Cells were treated with Scr (**I**), *Ppp3cb* (**J**), or *Ppp3r1* (**K**) siRNA for 48 h prior to infection with *S. aureus* (MOI = 10, t = 3 h). **L.** GFP-TFEB N/C Ratio by CellProfiler (3 biological replicates, n = 210 cells). *** p ≤ 0.001 (one-way ANOVA followed by Tukey’s post-hoc test). **M, N.** GFP-TFEB iBMDMs were treated with DMSO (**M**, t = 6 h) or Ionomycin (**N**, 10 μM, t = 6 h). **O.** GFP-TFEB N/C Ratio by CellProfiler (3 biological replicates, n = 350 cells). **** p ≤ 0.0001 (one-way ANOVA followed by Tukey’s post-hoc test). **P-S.** GFP-TFEB iBMDMs were treated with scrambled (Scr, **P**) or siRNA against *Mcoln1* (**Q**) for 48 h. Cells were treated with Scr (**R**) or *Mcoln1* (**S**) siRNA for 48 h prior to infection with *S. aureus* (MOI = 10, t = 3 h). **T.** GFP-TFEB N/C Ratio by CellProfiler (3 biological replicates, n = 300 cells). *** p ≤ 0.001 (one-way ANOVA followed by Tukey’s post-hoc test). **U, V.** GFP-TFEB iBMDMs were treated with DMSO (**U**, t = 3 h) or ML-SA1 (**V**, 10 μM, t = 3 h). **W.** GFP-TFEB N/C Ratio by CellProfiler (3 biological replicates, n = 355 cells). **** p ≤ 0.0001 (two-sample two-sided *t* test).

Previous studies have identified TRPML1/MCOLN1 as a major Ca^2+^ export channel in lysosomes ^22^. Furthermore, TRPML1/MCOLN1 was shown to function upstream of calcineurin for TFEB activation in several human and mouse cell types during nutrient starvation, and TRPML1/MCOLN1 was required for TFEB activation by FcγR during phagocytosis of opsonized beads in murine macrophages ^7,21^. In our hands, silencing of *Mcoln1* completely prevented TFEB activation and lysosome induction by *S. aureus* (**Fig. 4P-T**) and *Salmonella* (**Fig. S5M-Q**). Conversely, treatment of uninfected macrophages with TRPML1/MCOLN1 agonist ML-SA1 ^23^ resulted in strong TFEB nuclear import (**Fig. 4U-W**). Thus, in macrophages TRPML1/MCOLN1 is necessary and sufficient for TFEB activation during infection. Together, these results suggested that the TRPML1/MCOLN1-calcineurin-TFEB pathway is functional in macrophages, and is necessary and sufficient for TFEB nuclear import during infection. However, the mechanism of TRPML1/MCOLN1 activation during infection remained unknown.

### CD38 functions upstream of TRPML1/MCOLN1 for TFEB activation

Because of its relevance to lysosomal storage disorders, in particular mucolipidosis Type IV, the regulation of TRPML1/MCOLN1 is of great interest ^22^. Recently, it was reported that nicotinic acid adenine dinucleotide phosphate (NAADP) can function as a TRPML1/MCOLN1 ligand to induce lysosomal Ca^2+^ export in several cell types ^24,25^. Therefore, we investigated whether NAADP might function to induce TFEB activation downstream of MCOLN1 during infection in macrophages.

Prior treatment with the potent NAADP antagonist Ned-19 ^26^ rendered macrophages unable to translocate TFEB into the nucleus during infection with *S. aureus* (**Fig. 5A-C, F**). Likewise, treatment with kuromanin or apigenin, which inhibit the NAADP-synthesizing enzyme CD38 ^27,28^, also prevented TFEB activation by *S. aureus* (**Fig. 5D-F**). We obtained essentially the same results with *Salmonella* (**Fig. S6A-F**). Moreover, silencing of *Cd38* prevented TFEB activation by *S. aureus* (**Fig. 5G-J**). Together, these data showed that CD38 and NAADP were required for TFEB activation during infection. Conversely, treatment of macrophages with the NAADP analog NAADP-AM ^29^ resulted in strong induction of TFEB nuclear import (**Fig. 5K, L, N**). Together, these observations showed that CD38 and its product NAADP are necessary and sufficient for TFEB activation by *S. aureus* in macrophages. In contrast, NAADP-AM-induced TFEB activation was abrogated by silencing *Mcoln1* (**Fig. 5M, N**), suggesting that CD38 and NAADP require TRPML1/MCOLN1 to induce TFEB translocation, consistent with their proposed roles upstream of TRPML1/MCOLN1.

**Figure 5.**
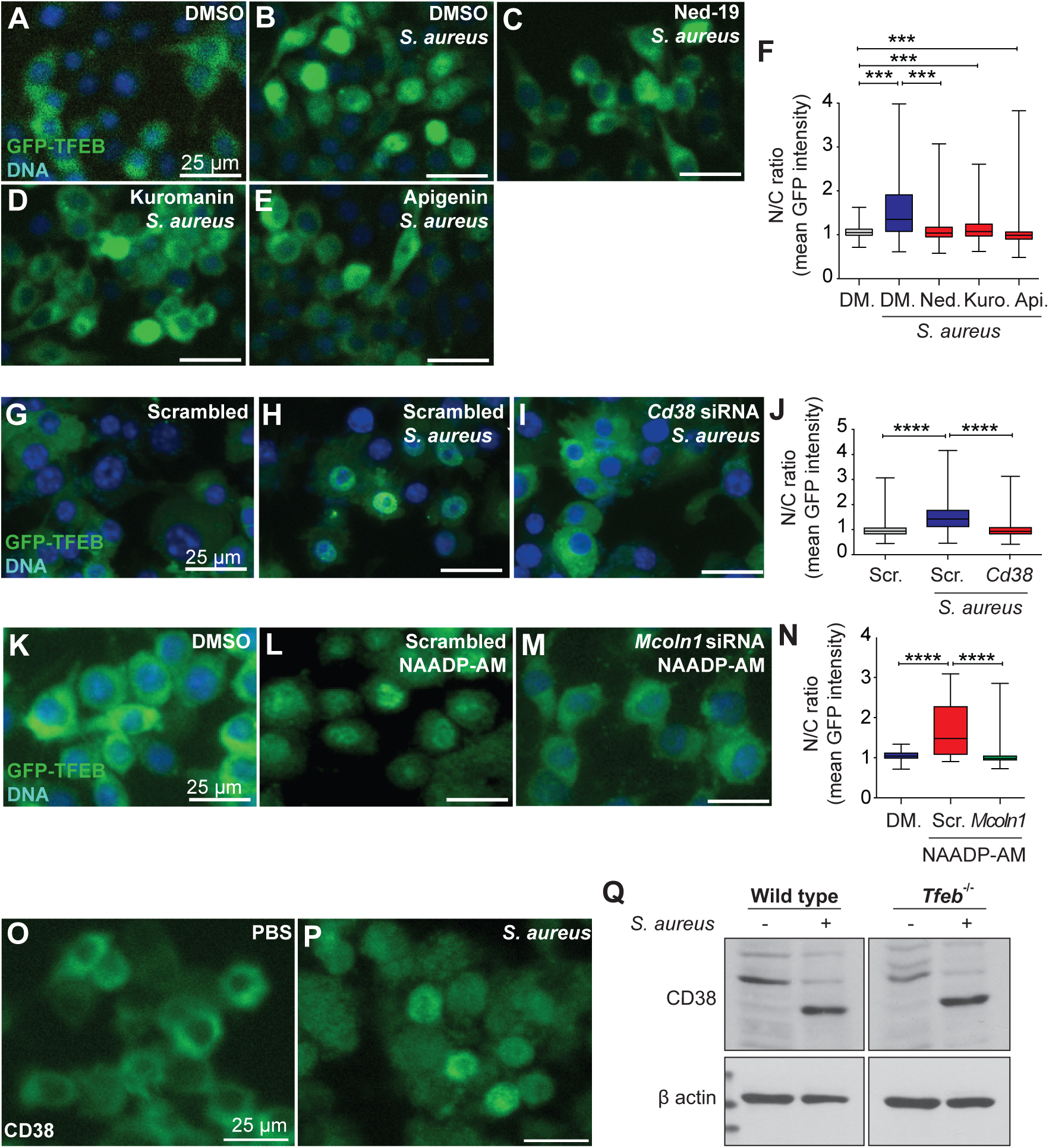
CD38 activates TFEB through NAADP and TRPML1/MCOLN1. **A-E.** GFP-TFEB iBMDMs were treated with DMSO without infection (DM., t = 6 h, **A**) or treated for 6 h and subsequently infected with *S. aureus* (**B**, MOI = 10, t = 3 h). In parallel, cells were treated with Ned-19 (**C,** Ned., 10 μM, t = 4 h), Kuromanin (**D,** Kuro., 100 μM, t = 6 h), or Apigenin (**E,** Api., 100 μM, t = 6 h) and subsequently infected with *S. aureus* (MOI = 10, t = 3 h). **F.** Quantification of GFP-TFEB N/C Ratio by CellProfiler (3 biological replicates, n = 200 cells). **** p ≤ 0.0001 (one-way ANOVA followed by Tukey’s post-hoc test). **G-I.** GFP-TFEB iBMDMs were treated with scrambled (Scr, **G**) or siRNA for 48 h. Cells were treated with Scr (**H**) or *Cd38* (**I**) siRNA for 48 h prior to infection with *S. aureus* (MOI = 10, t = 3 h). **J.** GFP-TFEB N/C Ratio by CellProfiler (3 biological replicates, n = 350 cells). *** p ≤ 0.001 (one-way ANOVA followed by Tukey’s posthoc test). **K-M.** GFP-TFEB iBMDMs were treated with DMSO (DM., t = 2 h, **K**) or treated with scrambled siRNA (Scr, t = 48 h) and subsequently incubated with NAADP-AM (100 nM, t = 2 h, **L**). In parallel, cells were treated with *Mcoln1* siRNA (t = 48 h) and subsequently incubated with NAADP-AM (100 nM, t = 2 h, **M**). **N.** Quantification of GFP-TFEB N/C Ratio by CellProfiler (3 biological replicates, n = 300 cells). **** p ≤ 0.0001 (one-way ANOVA followed by Tukey’s post-hoc test). **O, P.** CD38 immunofluorescence in wild type primary BMDM. PBS control (**O**) and *S. aureus* infection (MOI = 10, t = 2 h, **P**). **Q.** CD38 immunoblot of whole cell lysates from wild type and *Tfeb*^ΔLysM^ (*Tfeb*^-/-^) iBMDMs incubated with PBS or infected with *S. aureus* (MOI = 10, t = 3 h). β-actin is loading control. Representative of three biological replicates.

Immunofluorescence in untreated macrophages showed that CD38 localized principally to the plasma membrane, as previously shown in several human lymphocyte cell types ^30,31^. In contrast, *S. aureus-*infected macrophages exhibited intracellular staining in what appeared to be cytosolic vesicles (**Fig. 5O, P**). Such vesicles are presumably phagosomes ^32^. We also noticed increased staining in infected cells compared to baseline. To further examine this, we performed immunoblots. In wild type macrophages, we observed a slight increase in CD38 expression, albeit of a smaller molecular weight form (45 kDa v 64 kDa), in infected cells compared to control (**Fig. 5Q**). This increased expression occurred in *Tfeb*^-/-^ cells as well (**Fig. 5Q**). Together, these observations suggested that infection caused the relocalization of CD38 to endomembranes, its increased expression, and its presumed proteolytic conversion to a smaller form in a TFEB-independent manner.

### ROS function upstream of CD38 and of TFEB and TFE3

CD38 is a fascinating transmembrane enzyme with complex membrane topology, subcellular localization, and regulation ^31^. Upstream regulation of CD38 is poorly understood. It is thought that CD38 can be activated by reactive oxygen species (ROS) by an unknown mechanism ^33,34^. Moreover, ROS activated TRPML1/MCOLN1 in HEK293 cells, presumably through CD38 ^35^. Therefore, we investigated the role of ROS in TFEB activation during infection.

First, we verified the accumulation of ROS and H_2_O_2_ during infection with *S. aureus* in wild type macrophages (**Fig. 6A, B**). To our surprise, prior treatment with ROS-scavenging compounds N-acetyl cysteine (NAC) or N-acetyl cysteine aldehyde (NACA) completely prevented TFEB nuclear translocation by *S. aureus* (**Fig. 6C-G**) and *Salmonella* (**Fig. S6G-K**), suggesting that ROS were essential for TFEB activation during infection. Conversely, treatment with CCCP, which uncouples the mitochondrial electron transport chain and elicits ROS, caused TFEB translocation in the absence of infection (**Fig. 6H, I, K**). Silencing of *Cd38* completely abrogated TFEB translocation caused by CCCP (**Fig. 6J, K**). Together, these data indicated that ROS, generated during infection with *S. aureus*, are necessary and sufficient for TFEB activation by infection, in a mechanism that requires CD38. However, the source of ROS during phagocytosis remained unclear.

**Figure 6.**
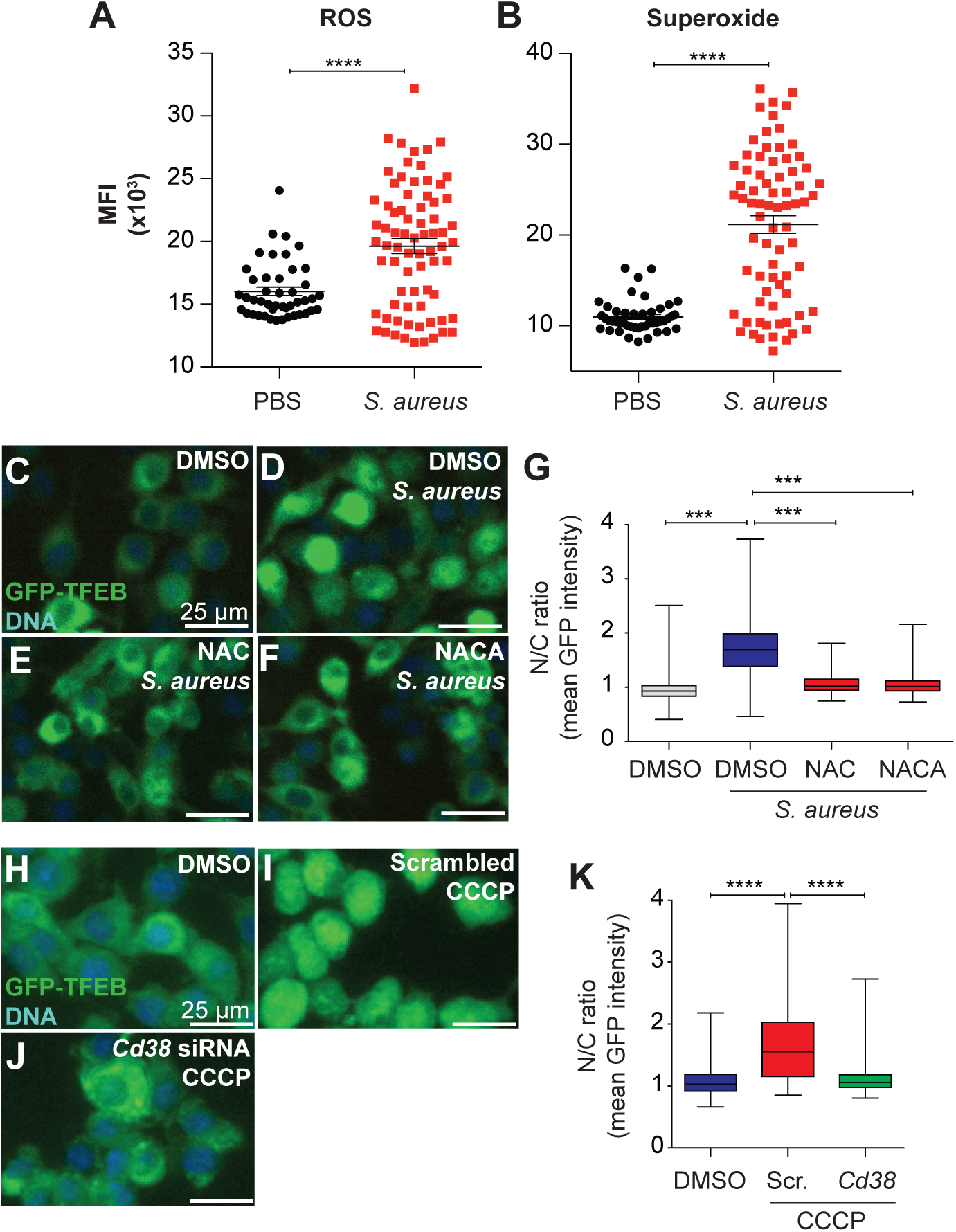
ROS activate TFEB through CD38. ROS (**A**) and superoxide (**B**) generated in wild type primary BMDM during infection with *S. aureus* (MOI = 10, t = 1 h) shown as mean fluorescence intensity (MFI) per cell and measured with Gen5 (n = 50 cells, 3 biological replicates). **** p ≤ 0.0001 (two-sided two-sample *t* test). **C-F.** GFP-TFEB iBMDMs were treated with DMSO without infection (DM., t = 4 h, **C**) or treated for 4 h and subsequently infected with *S. aureus* (**D**, MOI = 10, t = 3 h). In parallel, cells were treated with NAC (**E,** 5 mM, t = 4 h) or NACA (**F,** 1 mM, t = 4 h) and subsequently infected with *S. aureus* (MOI = 10, t = 3 h). **G.** Quantification of GFP-TFEB N/C Ratio by CellProfiler (3 biological replicates, n = 250 cells). *** p ≤ 0.001, **** p ≤ 0.0001 (one-way ANOVA followed by Tukey’s post-hoc test). **H-J.** GFP-TFEB iBMDMs were treated with DMSO (DM., t = 3 h, **H**) or treated with scrambled siRNA (Scr, t = 48 h) and subsequently incubated with CCCP (10 μM, t = 3 h, **I**). In parallel, cells were treated with *Cd38* siRNA (t = 48 h) and subsequently incubated with CCCP (10 μM, t = 3 h, **J**). **K.** Quantification of GFP-TFEB N/C Ratio by CellProfiler (3 biological replicates, n = 355 cells). **** p ≤ 0.0001 (one-way ANOVA followed by Tukey’s post-hoc test).

### NADPH oxidase functions upstream of the ROS-CD38-TRPML1/MCOLN1-CN-TFEB axis

During phagocytosis, recruitment of NADPH oxidase (PHOX) to the maturing phagosome is a key event for phagolysosome maturation and bactericidal killing^36^. PHOX catalyzes the production of H_2_O_2_ from NADPH oxidation, thus initiating the production of ROS in the phagolysosome ^37^. Therefore, we examined PHOX as a potential source of ROS for CD38 activation during bacterial phagocytosis.

First, we examined the oxidative burst in macrophages harboring a deletion in *Nox2,* which encodes the gp91phox subunit. While wild type macrophages infected with *S. aureus* induced high levels of ROS and superoxide, *Nox2*^-/-^ cells did not (**Fig. 7A, B**), consistent with PHOX being the main source of ROS in our infection assays. Consistent with the requirement for ROS to activate TFEB during infection, *S. aureus* did not induce TFEB nuclear translocation in *Nox2-* deficient cells (**Fig. 7C-G**), suggesting that PHOX is required for TFEB activation by infection. In support of this conclusion, chemical inhibition of PHOX activity using apocynin completely prevented TFEB activation during *S. aureus* infection (**Fig. H-K**). We observed similar results with *Salmonella* (**Fig. S6L-P**). Together, these data suggested that PHOX functions upstream of TFEB, and that during bacterial infection its activity is essential for TFEB activation.

**Figure 7.**
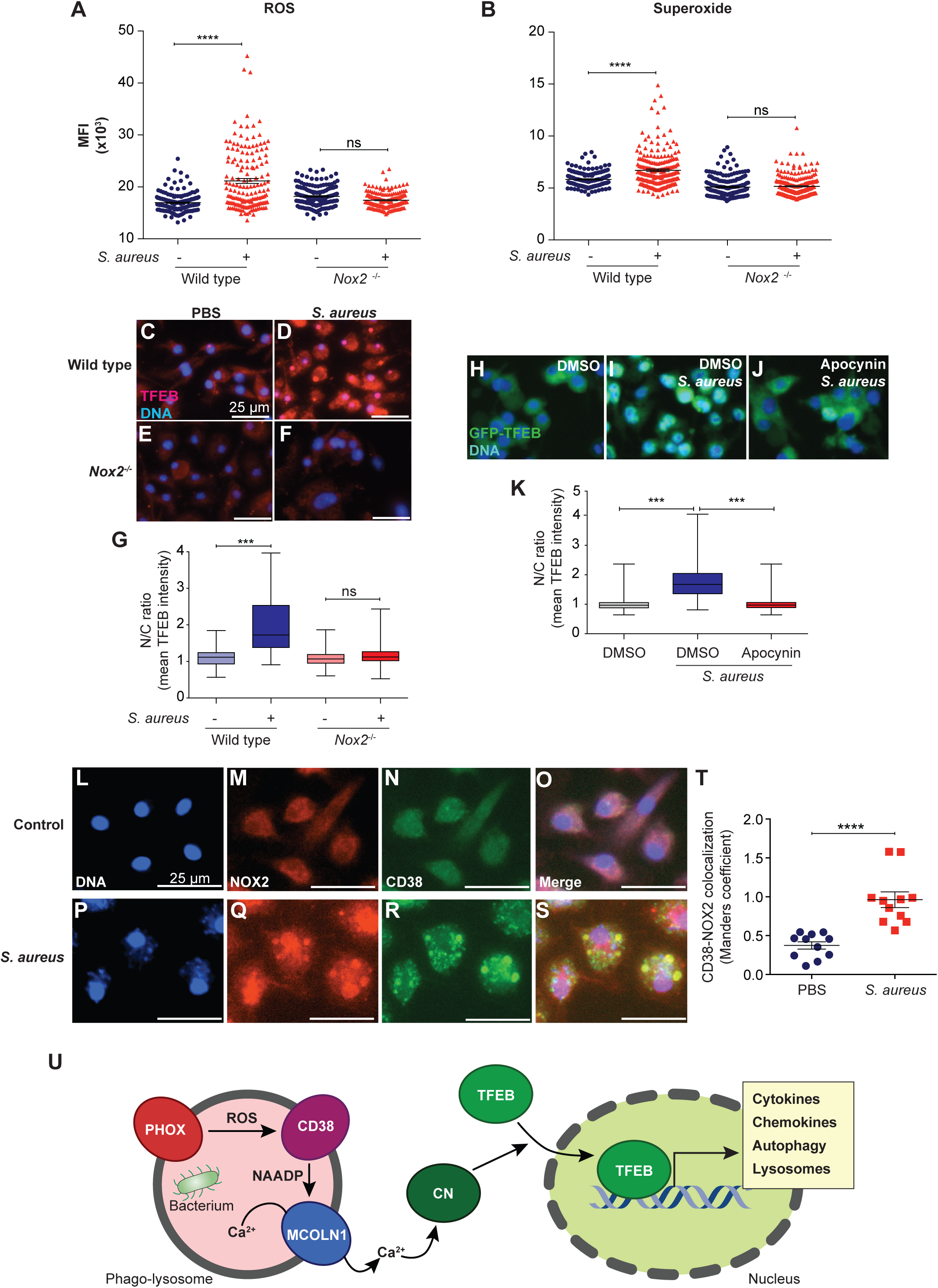
Infection induced ROS and TFEB activation require NADPH oxidase. ROS (**A**) and superoxide (**B**) generated in wild type and *Nox2*^-/-^ primary BMDM during infection with *S. aureus* (MOI = 10, t = 1 h) shown as mean fluorescence intensity (MFI) per cell and measured with Gen5 (n = 200 cells, 3 biological replicates). **** p ≤ 0.0001 (two-sided two-sample *t* test). **C-F.** TFEB immunofluorescence in wild type and *Nox2*^-/-^ primary BMDM. PBS controls (**C, E**) and infected cells (MOI = 10, t = 3 h, **D, F**). **G.** Quantification of TFEB N/C Ratio by CellProfiler (3 biological replicates, n = 200 cells). *** p ≤ 0.001, ns: p = 0.4399 (one-way ANOVA followed by Tukey’s post-hoc test). **H-J.** GFP-TFEB iBMDMs were treated with DMSO without infection (DM., t = 4 h, **H**) or treated for 4 h and subsequently infected with *S. aureus* (**I**, MOI = 10, t = 3 h). In parallel, cells were treated with Apocynin (**J,** 10 μM, t = 4 h) and subsequently infected with *S. aureus* (MOI = 10, t = 3 h). **K.** Quantification of GFP-TFEB N/C Ratio by CellProfiler (3 biological replicates, n = 250 cells). *** p ≤ 0.001 (one-way ANOVA followed by Tukey’s post-hoc test). **L-S.** Immunofluorescence detection of NOX2 and CD38 in wild type primary BMDM. Cells were incubated with PBS (**L-O**) or infected with *S. aureus* (MOI = 10, t = 3 h, **P-S**). **T.** Colocalization analysis of NOX2 and CD38. **** p ≤ 0.0001 (two-sided two-sample unpaired *t* test). **U**. Proposed hypothetical model for the phagocytosis-triggered activation of TFEB.

Immunofluorescence analysis revealed that *S. aureus* infection caused NOX2 and CD38 colocalization to increase about 100% over baseline (**Fig. 7L-T**). We obtained similar results with *Salmonella* (**Fig. S6Q-T**). Taken together, this series of experiments supports a hypothetical model (**Fig. 7U**) in which phagocytosis of bacterial cells may cause activated PHOX and CD38 to localize to the same compartment, likely the phago-lysosome. ROS produced by PHOX may cause CD38 to produce NAADP, which could activate TRPML1/MCOLN1 and induce Ca^2+^ export from the phago-lysosome/lysosome compartment. Ca^2+^ thus released may activate calcineurin, which can dephosphorylate TFEB. Dephosphorylated TFEB can translocate into the nucleus, where it drives the expression of cytokines and chemokines, as previously demonstrated ^4,6^. In addition, TFEB activation results in enhanced autophagy and lysosomal biogenesis, thus amplifying the phagocytic capacity of the cell via a positive feedback loop, and increases the degradative capacity of the cell via a positive feedforward loop ^7^. However, this model did not account for the requirement of the PC-PLC/PrKD1 pathway for TFEB activation.

### Disruption of PC-PLC/PrKD1 signaling inhibits TRPML1/MCOLN1 localization to the lysosome

A previous study showed that PrKD1 is required for TRPML1/MCOLN1 transport from the Golgi apparatus to the lysosome in HeLa cells and human skin fibroblasts ^38^. We tested if this is also a functional mechanism in macrophages. Using immunofluorescence in baseline murine macrophages, we observed low expression and localization to LAMP1-positive vesicles of TRPML1/MCOLN1 (**Fig. S7A-C**). TRPML1/MCOLN1 expression was highly upregulated by treatment with ML-SA1 and by infection with *S. aureus* (**Fig. SD-I, T**). Such upregulation was expected, since *Mcoln1* is a direct TFEB target in macrophages, and is upregulated by TFEB activation ^6^. Under *S. aureus* infection and MLSA1 treatment, we also observed higher colocalization of TRPML1/MCOLN1 and LAMP1 (**Fig. S7D-F, J-L, S**). *S. aureus* infection caused a 100% increase in TRPML1/MCOLN1 – LAMP1 colocalization, which was partially abrogated by prior treatment with PrKD1 inhibitor kb-NB142-70 (**Fig. S7M-O, S**). kb-NB142-70 also prevented the induced expression of TRPML1/MCOLN1 by *S. aureus* and MLSA1 (**Fig. S7T**). Together, these observations support the notion that PrKD1 is necessary for TFEB activation because it is required for TRPML1/MCOLN1 transport to the phagolysosome/lysosome compartment, where it is required for Ca^2+^ export downstream of NAADP.

## Discussion

In this article, we show a novel axis for TFEB activation in macrophages during phagocytosis of bacterial cells, which is essential for macrophage activation. Our observations support a model for a novel mechanism of pathogen sensing that is TLR-independent and is based on the phagocytic pathway. This pathway connects productive phagocytosis, presumably sensed by the recruitment of PHOX to the maturing phagosome. PHOX recruitment thus places the source of ROS in the same compartment as CD38, which may be endocytosed during phagocytosis of the bacterial particle. ROS generated by PHOX may activate CD38, as reported for lymphokine-activated killer cells and coronary arterial myocytes ^39,40^, causing the accumulation of NAADP in the phagosome. After phago-lysosome fusion, TRPML1/MCOLN1 may come into contact with this reservoir of NAADP, thus triggering the release of lysosomal Ca^2+^ into the cytosol. In the cytosol, Ca^2+^ activates calcineurin, which dephosphorylates TFEB and enables its nuclear import. In the nucleus, TFEB drives the expression of a large majority of genes that are induced upon pathogen phagocytosis. Such gene induction may require direct and indirect TFEB-dependent mechanisms. As far as we can tell, TFE3 is regulated in a similar manner.

Likewise, we found that bacterial molecules activate TFE3 and TFEB through a TLR-independent mechanism. These observations and the hypothetical model they support provide an explanation for previous, intriguing observations that phagocytosis is sufficient to trigger TFEB nuclear translocation, but only if the phagocytic cup is able to close ^7^. In this model, ROS serve dual functions as a bactericidal agent in the phago-lysosome, and as a signal to activate CD38 and initiate the signal cascade that results in TFEB/TFE3 activation. Since both PHOX- and mitochondria-derived ROS activated TFEB and TFE3, this provides redundant mechanisms of ROS production that report on phagocytosis and mitochondrial integrity. Thus, the model predicts that any condition that leads to PHOX activation or mitochondrial disruption, increasing ROS, would result in TFEB/3 activation (at least, in macrophages). This has important implications for homeostasis and disease.

Since our discovery of the key role of TFEB in the transcriptional host response to infection in macrophages ^4^, many studies have independently shown that TFEB and/or TFE3 greatly contribute to innate immunity against a large number of bacterial pathogens ^13,15,41^. In addition, several signaling components have been identified in the regulation of TFEB/3, including AGS3 during LPS stimulation, the activation of TFEB by IFN-γ, and the contribution of AMPK/mTORC1 regulation by several mechanisms ^42,43^. Moreover, at least one example of bacterial modulation of TFEB has been elucidated, particularly how *Mycobacterium tuberculosis* represses TFEB through *mir-33* and *mir-33\** ^44^. These discoveries underscore the central importance of TFEB/TFE3 as key contributors to the host response to infection, and provide a strong rationale for complete understanding of the contributions of the novel PHOX/CD38/MCOLN1/TFEB pathway described herein in distinct infectious diseases.

The events that occur downstream of TFEB/3 activation are of considerable interest. Prior studies demonstrated that TFEB and TFE3 have redundant and non-redundant functions important for the induction of key immune signaling molecules, such as intracellular signaling pathway components and extracellular cytokines and chemokines in LPS-stimulated macrophages ^6^. Here we show that this is also true in the case of bacterial phagocytosis. Thus, TFEB/3 are predicted to be key contributors to the function of cellular networks in tissues that are under pathogenic attack, both in the induction of antimicrobial responses and in the recruitment of effector cells of the innate and adaptive immune systems.

Furthermore, we show that TFEB activation in macrophages induces lysosomal biogenesis, including the induction of TRPML1/MCOLN1 expression. Enhanced lysosomal biogenesis increases the availability of TRPML1/MCOLN1-containing lysosomes for fusion with PHOX/CD38-positive maturing phagosomes, thus enacting a positive phagocytosis feedback loop. In addition, prior studies showed how TFEB activation in macrophages leads to increased autophagy ^45–47^, consistent with results in many other cell types. Thus, activation of the PHOX/CD38/TRPML1/CN/TFEB axis drives a positive phagocytosis feedforward loop that enhances the compartment that is poised to receive the mature phagolysosome for degradation, and enhances lysosomal exocytosis of remaining bacterial debris.

In summary, our studies reveal a heretofore unknown mechanism of macrophage activation that is activated by phagocytosis of bacterial cells. Such mechanism is parallel to better-understood TLR-mediated pathogen detection mechanisms. A key challenge for the future is to understand how diverse pathogen-detection pathways interact within an integrated regulatory network to produce a cellular response that is tailored to each particular pathogenic challenge. Additionally, understanding how pathogens have evolved to circumvent this new axis may help understand microbial pathogenesis and reveal new mechanisms to safely manipulate it for therapeutic purposes.

## Materials and Methods

### Isolation and differentiation of Bone Marrow Derived macrophages (BMDMs)

Femurs and tibias from 8-12 weeks old mice were separated and cleaned. Bone marrow was flushed into 50 ml tubes under the sterile hood. Bone marrow was passed through the bone until the color of the bone turned white. After centrifugation at 1500 rpm for 5 minutes at 4 °C, cell pellets were resuspended and plated in BMDM media: DMEM (Fisher Scientific, MT10102CV), FBS 10% (Thermo Fisher, 16000069), AA 1% (Life Technologies, 15240-062), L-glutamine 1% (Life Technologies, 25030-081), MEN NEA AA 1% (Life Technologies, 11140-050), 2-Mercapto 0.1% (Life Technologies, 21985), IL-3 5 ng/ml (Peprotech: 213-13), MCSF 5 ng/ml (Peprotech: 315-02). Cell were used for experiments after 1 week of differentiation. To produce immortalized BMDMs, cells were transformed by CreJ2 virus ^48^.

### Cell culture, transfection and Imaging

RAW264.7 and iBMDM cells were grown in Dulbecco’s Modified Eagle’s Medium (DMEM) - high glucose (Sigma-Aldrich, D6429-500ML) containing 10% endotoxin tested FBS (Thermo Fisher, 16000044), 1% Antibiotic-Antimycotic (Life Technologies 15240-062). Cells were passage 4 to 12 times. iBMDM GFP-TFEB stably transfected cells were created using pEGFP-N1-TFEB (pEGFP-N1-TFEB was a gift from Shawn Ferguson, Addgene plasmid # 38119), Lipofectamine 3000 (Thermo Fisher, L3000008) used according to manufacturer’s instructions, and G418 sulfate (Life Technologies, 10131) used for selection. 10 days after selection, stable cells were separated by FACS. For Imaging we used 96-Well Optical-Bottom Plates (Thermo Fisher, 165305). 6×10^4^ cells were seeded in each well. At the end of the experiments, cells were fixed using 4% paraformaldehyde (Sigma-Aldrich, 158127) and incubated with Hoechst stain (Anaspec, AS-83218) at room temperature for 20 minutes as nuclear staining. For lysotracker staining, LysoTracker Red DND-99 (Thermo Fisher, L7528) was added to the media 30 minutes before fixing according to the manufacture instruction. Image acquisition was automatically performed using a Cytation3 Imaging Plate Reader (Biotek). The N/C ratio was measured using CellProfiler (Broad Institute), as described in^5^. Colocalization analysis was performed using ImageJ software (NIH).

### siRNA Knockdown

siRNA compounds were purchased from Dharmacon RNAi Technologies. siGENOME NonTargeting siRNA #1 (D-001210-01-05). siGENOME Mouse Cd38 (12494) siRNA - SMARTpool (M-058632-01-0005). siGENOME Mouse Ppp3cb (19056) siRNA (M-063545-00-0005). siGENOME Mouse Ppp3r1 (19058) siRNA (M-040744-01-0005). siGENOME Mouse Mcoln1 (94178) siRNA (M-044469-00-0005). We used Lipofectamine RNAiMAX (Thermo Scientific, 13778030) for transfection according to manufacturer’s instructions.

### Bacterial strains

*Salmonella enterica* serovar Typhimurium SL1344 wild type is a gift from Brian Coombes (McMaster University, Canada), *Staphylococcus aureus* NCTC8325 is a gift from Fred Ausubel, MGH Research Institute, USA.

### In vitro infection

Bacteria were grown overnight at 37 °C in LB medium (Difco, BD) with 100 μg/ml streptomycin for Salmonella and Columbia medium (Difco, BD) with 10 μg/ml Nalidixic acid for *S. aureus*. The following day cultures were diluted 1:50 in the same medium and grown at 37 °C for 3 h to late-exponential phase, washed twice in cold PBS, and cells were infected with MOI 10 for *S. aureus* and MOI 10 for S*. enterica*, as in (Engelenburg and Palmer, 2010; Najibi et al., 2016; Trieu et al., 2009; Visvikis et al., 2014). For gentamycin antibiotic (AB) - killed bacteria, before addition to cells, gentamicin (100 µg/ml) was added to washed bacteria in PBS for 2 hours and 100% killing of the bacteria was confirmed by culture for 48 h on LB-streptomycin agar at 37°C. The appropriate amounts of bacteria were resuspended in DMEM 10 % FBS without antibiotic.

### Immunofluorescence

After treatment, cells were fixed with 4% paraformaldehyde (PFA) pH 7.4 at room temperature for 10 min and washed 3 times in PBS (Gibco Life Technologies,10010) for 5 min each. PFA was neutralized with 50 mM NH_4_Cl in PBS at room temperature for 10 minutes with agitation. After 3 washes with PBS, cells were permeabilized with 0.1% Triton X in PBS at room temperature on agitator for 5 min and then blocked with 5% bovine serum albumin (Sigma-Aldrich, A9647) in PBS for 1 h. After 3 washes with PBS cells were incubated with primary antibodies for 2 hours. Anti-TFEB Antibody (Cell Signaling, 4240). Anti-TFE3 antibody (Sigma-Aldrich, HPA023881-100UL). Anti-CD38 antibody (Abcam, ab24978). Anti-NOX2/gp91phox antibody (Abcam, ab80508). Anti-LAMP1 (Cell Signaling Technology, 9091). Anti-Mucolipin 1 (MCOLN1) (C-Term) antibody (Antibodies-Online, ABIN571446). Cells were washed thrice in PBS and incubated with the fluorescent secondary antibody plus Hoechst stain (Anaspec, AS-83218) at room temperature for 1 h. Image acquisition was performed using a Cytation3 imaging plate reader (Biotek).

### Immunoblotting

Cells were washed 3 times with PBS, harvested, and lysed with 1X SDS sample buffer Blue Loading Pack (Cell Signaling, 7722) at 100 µl per well of 6-well plate. Lysates were heated at 100 °C for 5 min and then centrifuged for 5 min. The supernatant was collected and sonicated, gel electrophoresis was performed using Novex 4-20% Tris-Glycine Mini Gels (Invitrogen, XP04200BOX), and were then transferred onto nitrocellulose (Life Technologies, LC2009). After wash with TBS (Life Technologies, 28358) for 5 minutes, membranes were soaked in blocking buffer containing 1X TBS with 5% BSA for 1 hour at room temperature. After 3 washes with TBS-Tween (Life Technologies, 28360), membranes were incubated overnight at 4 °C with primary antibodies and gentle agitation. Next membranes were washed thrice with TBS-Tween and incubated with HRP-conjugated secondary antibody (Cell Signaling, 7074 1:2000) for 1 h at room temperature with gentle agitation. Membranes were then washed with TBS-Tween and incubated with LumiGLO® (Cell signaling, 7003) for 1 min and exposed to x-ray film (Denville Scientific, E3012). Primary antibodies and dilutions that were used are as follows: β-actin antibody (Cell Signaling Technology, 4967, 1:1000), Anti-CD38 Antibody (Cell Signaling Technology, 14637S 1:1000), Anti-TLR4 antibody (). Anti-TLR2 antibody (). Anti-Mucolipin 1 (MCOLN1) (C-Term) antibody (Antibodies-Online, ABIN571446 1:1000).

### Drugs and reagents

Lipopolysaccharides (LPS) from *S. enterica* serotype Typhimurium (Sigma-Aldrich, L6143-1MG, 10ug/ml): Peptidoglycan (PGN) from *S. aureus* (Invivogen, tlrl-pgns2): Monophosphoryl Lipid A (MPLA-SM) (Invivogen, tlrl-mpla): ML-SA1 (Sigma Aldrich, SML0627) Ionomycin (Cayman Chemical Company, 10004974): FK-506 (Cayman Chemical Company, 10007965): BAPTA AM (Cayman Chemical Company, 15551): Ned-19 (Cayman Chemical Company, 17527): Kuromanin (Cayman Chemical Company, 16406): Apigenin (Cayman Chemical Company, 10010275): CCCP (Cayman Chemical Company, 25458): N-acetyl-L-Cysteine (NAC) (Cayman Chemical Company, 20261): N-acetyl-L-Cysteine amide (NACA) (Cayman Chemical Company, 25866): Apocynin (Cayman Chemical Company, 11976): D609 (Cayman Chemical, 13307, 50 µM): kb NB 142-70 (Cayman Chemical Company, 18002).

### ROS/Superoxide Detection Assay

Cellular ROS/Superoxide Detection Assay Kit (Abcam, ab139476) was used according to the manufacture instruction. Mean fluorescence intensity was calculated by Biotek Gen5 Data Analysis Software.

### RNA Sequencing and Differential Expression Analysis

RAW264.7 dKO were a one-time gift from Rosa Puertollano (NHLBI). Infection was performed as described in *In Vitro Infection,* in three independent replicates. RNA was extracted by TRIzol Plus RNA Purification Kit (Thermo Fisher, 12183555). Whole cell RNA was submitted to BGI Inc for ribosomal RNA depletion, library construction, and sequencing on Illumina NextSeq machines. Reads were trimmed and quality-filtered by BGI. Clean reads were mapped to the GRCm38 mouse reference transcriptome using Salmon v0.13.1 and transcript-quantified (Patro et al., 2017). Salmon quant outputs were analyzed using tximport and DESeq2 v1.22.2 in R (Love et al., 2014). DESeq2 output counts were used as input for interactive analysis and data plotting using DEBrowser 1.10.9 in R (Kucukural et al., 2019). Differentially expressed (fold change ≥ 2; Padj ≤ 0.01; filter genes with fewer than 10 counts) gene sets were used as input for g:Profiler v0.6.6 online tools (Raudvere et al., 2019).

### ELISA Assay

Concentrations of IL-1α, IL-1β, IL-6 and TNF-α were measured by enzyme-linked immunosorbent assays (ELISAs) using kits purchased from R&D Systems (DY400-05, DY401-05, DY406-05, DY8234-05, DY410-05, respectively). All incubation periods occurred at room temperature, and plates were washed with Tris buffered saline (TBS) containing 0.05% Tween 20 (Bethyl Laboratories, E106) after each step. Briefly, clear microplates (R&D Systems, DY990) were incubated overnight with capture antibody. Plates were blocked for one hour using TBS containing 1% BSA (Bethyl Laboratories, E104). Samples were added undiluted (IL-1α, IL-1β, IL-6) or diluted 1:10 in TBS containing 1% BSA (TNF-α) and plates were incubated for two hours. Detection antibody was added, and plates were incubated for two hours. Streptavidin-Horseradish Peroxidase (Strep-HRP) was added, and plates were incubated for 20 minutes. TMB One Component HRP Microwell Substrate (Bethyl Laboratories, E102) was added, and reaction was stopped after 20 minutes using ELISA Stop Solution (Bethyl Laboratories, E115). Optical densities were measured at 450 nm and 540 nm using SpectraMax Plus 384 Microplate Reader with SoftMax Pro software. Measurements at 540 nm were subtracted from measurements at 450 nm to correct for optical impurities of the plates, per manufacturer’s recommendations. Concentrations of samples were interpolated from each standard curve using Prism 8. Data are represented as mean ± SEM.

### LDH Assay

Lactate Dehydrogenase (LDH) activity was measured using an LDH assay kit purchased from Abcam (ab102526). LDH assay was performed following manufacturer’s guidelines. Briefly, samples were added to a clear microplate (R&D Systems, DY990). Reaction mix was made using supplied LDH Assay Buffer and LDH Substrate Mix. Reaction mix was then added to each sample. Optical densities were measured at 450 nm using SpectraMax Plus 384 Microplate Reader with SoftMax Pro software. The amount of NADH in each sample was interpolated from the standard curve using Prism 8. LDH activity was quantified by dividing the amount of NADH in each sample by the product of the reaction time in minutes and the original volume of sample added into the reaction well in mL. LDH activity was reported in mU/mL and plotted as mean ± SEM.

## Acknowledgments

The authors are grateful to members of the Irazoqui laboratory, the Fitzgerald laboratory, the Program in Innate Immunity, and the Department of Microbiology and Physiological Systems for helpful insights and discussions. The Fitzgerald laboratory provided several knockout primary and immortalized BMDMs, and the Sassetti laboratory provided *Nox2* mutant mice and CreJ2 virus. This work was partially funded by grant R01GM101056 (JEI) from the National Institutes of Health. The content is solely the responsibility of the authors and does not necessarily represent the official views of the National Institutes of Health.

## Author Contributions

Conceptualization, JEI and MN; Methodology, JEI and MN; Investigation, MN, JAM, JEI, and HHH; Animals, JAM; Writing–original draft, JEI; Writing–review and editing, JEI, MN, JAM, and HH; Visualization, MN and JEI; Supervision, JEI; Funding acquisition, JEI.

