## Supplemental Text for "A Novel PHOX/CD38/MCOLN1/TFEB Axis Important For Macrophage Activation During Bacterial Phagocytosis"

**Figure S1. Gram- bacterial cells and ligands activate TFE3. A-H.** TFEB-GFP RAW264.7 cells were incubated with *Salmonella* Typhimurium (live *Salmonella*, MOI = 10), Antibiotic killed *Salmonella* Typhimurium (dead *Salmonella*, MOI = 10) and *Salmonella* Typhimurium lipopolysaccharide (LPS, 1 µg/ml) for different time points from 0 to 360 minutes after incubation. Cells were fixed and processed for anti-TFE3 immunofluorescence staining. Shown are representative images, and quantification of three biological replicates. Fluorescence intensity in the nucleus compared to the cytoplasm (N/C ratio) measured using CellProfiler. **** p ≤ 0.0001 (one-way ANOVA followed by Tukey’s post-hoc test (n = 400). **A** Images from anti-TFE3 immunofluorescence at 0, 120 and 320 minutes after *Salmonella* Typhimurium (live *Salmonella*, MOI = 10). **B**. Images from anti-TFE3 immunofluorescence at 0, 120 and 320 minutes after Antibiotic killed *Salmonella* Typhimurium (dead *Salmonella*, MOI = 10), and **C**. TFE3 N/C ratio quantification for *Salmonella* Typhimurium (live *Salmonella*, MOI = 100). **D**. TFE3 N/C ratio quantification for Antibiotic killed *Salmonella* Typhimurium (dead *Salmonella*, MOI = 100), **E**. TFE3 N/C ratio quantification for *Salmonella* Typhimurium lipopolysaccharide (LPS, 1 µg/ml)*,* **F**. TFEB N/C ratio quantification for *Salmonella* Typhimurium (live *Salmonella*, MOI = 100). **G**. TFEB N/C ratio quantification for Antibiotic killed *Salmonella* Typhimurium (dead *Salmonella*, MOI = 100), **H**. TFEB N/C ratio quantification for *Salmonella* Typhimurium lipopolysaccharide (LPS, 1 µg/ml).

**Figure S2. Gram- bacterial cells and ligands activate TFE3*.*** **A-C**. BMDMs from *Myd88-Trif* double knockout mice (*Myd88-Trif* dKO) and wild type mice were used for these experiments. *S. aureus* (MOI = 10) for 3 h. *Salmonella* Typhimurium (MOI = 1) for 6 h. Incubation with PGN from *S. aureus* was done for 3 h. Incubation with *Salmonella* Typhimurium lipopolysaccharide (LPS, 1 µg/ml) was done for 3 h. Lysotracker staining (50nM) was done for 1 h and cells were fixed. ) Lysotracker intensity (MFI/cell) in each cell measured by CellProfiler. **** p ≤ 0.0001 (one-way ANOVA followed by Tukey’s post-hoc test). Shown are representative images, and quantification of three biological replicates. **A.** Lysotracker staining in wild type BMDMs. **B.** Lysotracker staining in *Myd88-Trif* dKO BMDMs. **C.** Lysotracker intensity (MFI/cell) in each cell.

**Figure S3. TLR signaling is dispensable for TFEB activation. A-X**. iBMDMs from wild type (WT), TLR2 knockout (TLR2KO) and TLR4 knockout (TLR4KO) were used for the experiments. Cells were fixed and immunofluorescence staining was done with an antibody against TFEB. Shown are representative images from one replicate, and quantification of three biological replicates. TFEB fluorescence intensity in the nucleus compared to the cytoplasm (N/C ratio) was measured using CellProfiler. **** p ≤ 0.0001 (one-way ANOVA followed by Tukey’s post-hoc test). **A**. Immunoblot against TLR2 and β actin in TLR2 knockout (TLR2KO) cells. **B.** Immunoblot against TLR4 and β actin in TLR4 knockout (TLR4KO) cells. **C**. DMSO control in WT cells. **D**. *S. aureus* peptidoglycan (PGN, 10 µg/ml, t = 3 h) in WT cells. **E**. pam3csk4 (10 µg/ml, t = 3 h) in WT cells. **F**. *S. aureus* (Sa, MOI = 10, t = 3 h) in WT cells. **G**. Starvation (t = 3 h) in WT cells. **H**. DMSO control in TLR2KO cells. **I**. *S. aureus* peptidoglycan (PGN, 10 µg/ml, t = 3 h) in TLR2KO cells. **J**. pam3csk4 (10 µg/ml, t = 3 h) in TLR2KO cells. **K**. *S. aureus* (Sa, MOI = 10, t = 3 h) in TLR2KO cells. **L**. Starvation (t = 3 h) in TLR2KO cells. **M**. DMSO control in WT cells. **N**. *Salmonella* Typhimurium (*Salmonella,* MOI = 1, t = 6 h) in WT cells. **O.** *Salmonella* Typhimurium lipopolysaccharide (LPS, 1 µg/ml, t = 6 h) in WT cells. **P**. Monophosphoryl lipid A (MPLA, 1 µg/ml, t = 6 h) in WT cells. **Q**. Starvation (t = 3 h) in WT cells. **R**. DMSO control in TLR4KO cells. **S**. *Salmonella* Typhimurium (*Salmonella,* MOI = 1, t = 6 h) in TLR4KO cells. **T.** *Salmonella* Typhimurium lipopolysaccharide (LPS, 1 µg/ml, t = 6 h) in TLR4KO cells. **U**. Monophosphoryl lipid A (MPLA, 1 µg/ml, t = 6 h) in TLR4KO cells. **V**. Starvation (t = 3 h) in TLR4KO cells. **W**. TFEB fluorescence intensity in the nucleus compared to the cytoplasm (N/C ratio) in WT cells. **X**. TFEB fluorescence intensity in the nucleus compared to the cytoplasm (N/C ratio) in TLR4KO cells.

**Figure S4. Bacterial activation of TFE3 requires the PC-PLC/PrKD1 pathway. A-P.** Wild type BMDMs were used for these experiments. Cells were fixed and immunofluorescence staining was done with antibody against TFE3 and TFEB. Shown are representative images from one replicate, and quantification of three biological replicates. TFEB-GFP intensity in the nucleus compared to the cytoplasm (N/C ratio) was measured using CellProfiler. *** p ≤ 0.001 (one-way ANOVA followed by Tukey’s post-hoc test). **A.** Image from anti-TFE3 immunofluorescence, DMSO (t = 6 h). **B.** Image from anti-TFEB immunofluorescence, DMSO (t = 6 h). **C.** Image from anti-TFE3 immunofluorescence, *Salmonella* Typhimurium (*Salmonella*, MOI = 10, t = 6 h). **D.** Image from anti-TFEB immunofluorescence, *Salmonella* Typhimurium (*Salmonella*, MOI = 10, t = 6 h). **E.** Image from anti-TFE3 immunofluorescence, kb-NB142-70 (10 μM, t = 3 h) before infection with *Salmonella* Typhimurium (*Salmonella*, MOI = 10, t = 6 h). **F.** Image from anti-TFEB immunofluorescence, kb-NB142-70 (10 μM, t = 3 h) before infection with *Salmonella* Typhimurium (*Salmonella*, MOI = 10, t = 6 h). **G.** Image from anti-TFE3 immunofluorescence, tricyclodecan-9-yl-xanthogenate (D609, 50 μM, t = 3 h) before infection with *Salmonella* Typhimurium (*Salmonella*, MOI = 10, t = 6 h). **H.** Image from anti-TFE3 immunofluorescence, tricyclodecan-9-yl-xanthogenate (D609, 50 μM, t = 3 h) before infection with *Salmonella* Typhimurium (*Salmonella*, MOI = 10, t = 6 h). **I**. TFE3 N/C ratio quantification for *Salmonella* Typhimurium (*Salmonella*, MOI = 10, t = 6 h). **J**. TFE3 N/C ratio quantification for *Salmonella* Typhimurium lipopolysaccharide (LPS, 1 µg/ml, t = 6 h). **K**. TFE3 N/C ratio quantification for *S. aureus* (MOI = 10, t = 3 h). **L**. TFEB N/C ratio quantification for *Salmonella* Typhimurium (*Salmonella*, MOI = 10, t = 6 h). **M**. TFEB N/C ratio quantification for *Salmonella* Typhimurium lipopolysaccharide (LPS, 1 µg/ml, t = 6 h). **N**. TFEB N/C ratio quantification for *S. aureus* (MOI = 10, t = 3 h). **O**. TFE3 N/C ratio quantification for DAG analog, phorbol 12-myristate 13-acetate (PMA, 100ng/ml, t = 1 h). **P**. TFEB N/C ratio quantification for DAG analog, phorbol 12-myristate 13-acetate (PMA, 100ng/ml, t = 1 h).

**Figure S5. Bacterial activation of TFEB requires the TRPML1/MCOLN1 – Ca2+ – calcineurin pathway. (A-Q)** TFEB-GFP iBMDMs were used for all the experiments. Lysotracker staining (50 nM) was done for 1 h. Shown are representative images from one replicate, and quantification of three biological replicates. TFEB-GFP intensity in the nucleus compared to the cytoplasm (N/C ratio) was measured using CellProfiler. (**A**) DMSO control. (**B**) *Salmonella* Typhimurium (*Salmonella*, MOI = 10, t = 6 h)*.* (**C**) BAPTA (10 µM) was added 3 h before *Salmonella* Typhimurium (*Salmonella*, MOI = 10, t = 6 h). (**D**) FK506 (5 µM) was added 6 h before *Salmonella* Typhimurium (*Salmonella*, MOI = 10, t = 6 h). (**E**) TFEB-GFP intensity in the nucleus compared to the cytoplasm (N/C ratio). **** p ≤ 0.0001 (one-way ANOVA followed by Tukey’s post-hoc test). (**F**) scrambled siRNA (Scr) (**G**) siRNA against *Ppp3cb* (calcineurin subunit). (**H**) siRNA against *Ppp3r1* (calcineurin subunit). (**I**) scrambled siRNA (Scr) was performed 48 h before *Salmonella* Typhimurium (*Salmonella*, MOI = 10, t = 6 h). (**J**) siRNA against *Ppp3cb* was performed 48 h before *Salmonella* Typhimurium (*Salmonella*, MOI = 10, t = 6 h)*.* (**K**) siRNA against *Ppp3r1* was performed 48 h before *Salmonella* Typhimurium (*Salmonella*, MOI = 10, t = 6 h). (**L**) TFEB-GFP intensity in the nucleus compared to the cytoplasm (N/C ratio). **** p ≤ 0.0001 (one-way ANOVA followed by Tukey’s post-hoc test). (**M**) scrambled siRNA (Scr). (**N**) siRNA against *Mcoln1.* (**O**) scrambled siRNA (Scr) was performed 48 h before *Salmonella* Typhimurium (*Salmonella*, MOI = 10, t = 6 h). (**P**) siRNA against *Mcoln1* was performed 48 h before *Salmonella* Typhimurium (*Salmonella*, MOI = 10, t = 6 h). (**Q**) TFEB-GFP intensity in the nucleus compared to the cytoplasm (N/C ratio). **** p ≤ 0.0001 (one-way ANOVA followed by Tukey’s post-hoc test). (**U**) DMSO control.

**Figure S6. ROS activate TFEB through CD38.** (**A-F**) TFEB-GFP iBMDMs were used for the experiments. Shown are representative images from one replicate, and quantification of three biological replicates. TFEB-GFP intensity in the nucleus compared to the cytoplasm (N/C ratio) was measured using CellProfiler. *** p ≤ 0.001. (one-way ANOVA followed by Tukey’s post-hoc test). (**A**) DMSO control (DM). (**B**) *Salmonella* Typhimurium (*Salmonella*, MOI = 10, t = 6 h)*.* (**C**) Ned-19 (Ned) (10 µM) was added 4 h before *Salmonella* Typhimurium (*Salmonella*, MOI = 10, t = 6 h). (**D**) Kuromanin (Kuro) (100uM) was added 6 h before *Salmonella* Typhimurium (*Salmonella*, MOI = 10, t = 6 h). (**E**) Apigenin (Api) (100uM) was added 6 h before *Salmonella* Typhimurium (*Salmonella*, MOI = 10, t = 6 h). (**F**) TFEB-GFP intensity in the nucleus compared to the cytoplasm (N/C ratio). (**G-K)** GFP-TFEB iBMDMs were treated with DMSO without infection (DM., t = 4 h, **G**) or treated for 4 h and subsequently infected with *Salmonella* (**H**, MOI = 10, t = 6 h). In parallel, cells were treated with NAC (**I,** 5 mM, t = 4 h) or NACA (**J,** 1 mM, t = 4 h) and subsequently infected with *Salmonella* (MOI = 10, t = 6 h). **K.** Quantification of GFP-TFEB N/C Ratio by CellProfiler (3 biological replicates, n = 300 cells). *** p ≤ 0.001 (one-way ANOVA followed by Tukey’s post-hoc test). (**L-P**) BMDMs from wild type and *Nox2* knockout mice (*Nox2*^-/-^) were used for immunofluorescence against TFEB. (**L**) PBS control in wild type BMDMs. (**M**) *Salmonella* Typhimurium (*Salmonella*, MOI = 10, t = 6 h) in wild type BMDMs*.* (**N**) PBS control in *Nox2*^-/-^ BMDMs. (**O**) *Salmonella* Typhimurium (*Salmonella*, MOI = 10, t = 6 h) in *Nox2*^-/-^ BMDMs*.* (**P**) TFEB fluorescence intensity in the nucleus compared to the cytoplasm (N/C ratio), measured by CellProfiler. *** p ≤ 0.001. (one-way ANOVA followed by Tukey’s post-hoc test). (**Q-W**) Wild type BMDMs were incubated with PBS as control and *Salmonella* Typhimurium (*Salmonella*, MOI = 10, t = 6 h)*,* immunofluorescence against NOX2 and CD38 were done. Colocalization analysis was done using ImageJ software (NIH). **** p ≤ 0.0001. (two-sample t test). (**Q**) Immunofluorescence for NOX2, PBS control. (**R**) Immunofluorescence for CD38, PBS control. (**S**) Merge for NOX2 and CD38, PBS control. (**T**) Immunofluorescence for NOX2, *Salmonella* Typhimurium (*Salmonella*, MOI = 10, t = 6 h). (**U**) Immunofluorescence for CD38, *Salmonella* Typhimurium (*Salmonella*, MOI = 10, t = 6 h). (**V**) Merge for NOX2 and CD38, *Salmonella* Typhimurium (*Salmonella*, MOI = 10, t = 6 h). (**W**) Colocalization analysis for NOX2 and CD38, comparing the PBS control and *S. aureus* infection.

**Figure S7. Disruption of PC-PLC/PrKD1 signaling inhibits TRPML1/MCOLN1 localization to the lysosome.** (**A-T**) Wild type BMDMs were used for the experiments*.* Immunofluorescence against MCOLN1 and LAMP1 were done. Colocalization analysis was done using ImageJ software (NIH). **** p ≤ 0.0001, *** p ≤ 0.001. (one-way ANOVA followed by Tukey’s post-hoc test). MCOLN1 mean fluorescence intensity was measured by CellProfiler. **** p ≤ 0.0001, (one-way ANOVA followed by Tukey’s post-hoc test). (**A**) Immunofluorescence for MCOLN1, DMSO control (t = 6 h). (**B**) Immunofluorescence for LAMP1, DMSO control (t = 6 h). (**C**) Merge for MCOLN1 and LAMP1, DMSO control (t = 6 h). (**D**) Immunofluorescence for MCOLN1, ML SA1 (10 µM, t = 3 h). (**E**) Immunofluorescence for LAMP1, ML SA1 (10 µM, t = 3 h). (**F**) Merge for MCOLN1 and LAMP1, ML SA1 (10 µM, t = 3 h). (**G**) Immunofluorescence for MCOLN1, kb-NB142-70 (Kb, 10 μM, t = 3 h) before ML SA1 (10 µM, t = 3 h). (**H**) Immunofluorescence for LAMP1, kb-NB142-70 (Kb, 10 μM, t = 3 h) before ML SA1 (10 µM, t = 3 h). (**I**) Merge for MCOLN1 and LAMP1, kb-NB142-70 (Kb, 10 μM, t = 3 h) before ML SA1 (10 µM, t = 3 h). (**J**) Immunofluorescence for MCOLN1, *S. aureus* (MOI = 10, t = 3 h). (**K**) Immunofluorescence for LAMP1, *S. aureus* (MOI = 10, t = 3 h). (**L**) Merge for MCOLN1 and LAMP1, *S. aureus* (MOI = 10, t = 3 h). (**M**) Immunofluorescence for MCOLN1, kb-NB142-70 (Kb, 10 μM, t = 3 h) before *S. aureus* (MOI = 10, t = 3 h). (**N**) Immunofluorescence for LAMP1, kb-NB142-70 (Kb, 10 μM, t = 3 h) before *S. aureus* (MOI = 10, t = 3 h). (**O**) Merge for MCOLN1 and LAMP1, kb-NB142-70 (Kb, 10 μM, t = 3 h) before *S. aureus* (MOI = 10, t = 3 h). (**P**) Detail, Merge for MCOLN1 and LAMP1, DMSO control (t = 6 h). (**Q**) Detail, Merge for MCOLN1 and LAMP1, *S. aureus* (MOI = 10, t = 3 h). (**R**) Detail, Merge for MCOLN1 and LAMP1, kb-NB142-70 (Kb, 10 μM, t = 3 h) before *S. aureus* (MOI = 10, t = 3 h). (**S**) Colocalization analysis was done using ImageJ software (NIH). (**T**) MCOLN1 mean fluorescence intensity was measured by CellProfiler.
