## Supplementary figures and images for "A Novel PHOX/CD38/MCOLN1/TFEB Axis Important For Macrophage Activation During Bacterial Phagocytosis"

### Figure S1

**Najibi et al - FIGURE S1**

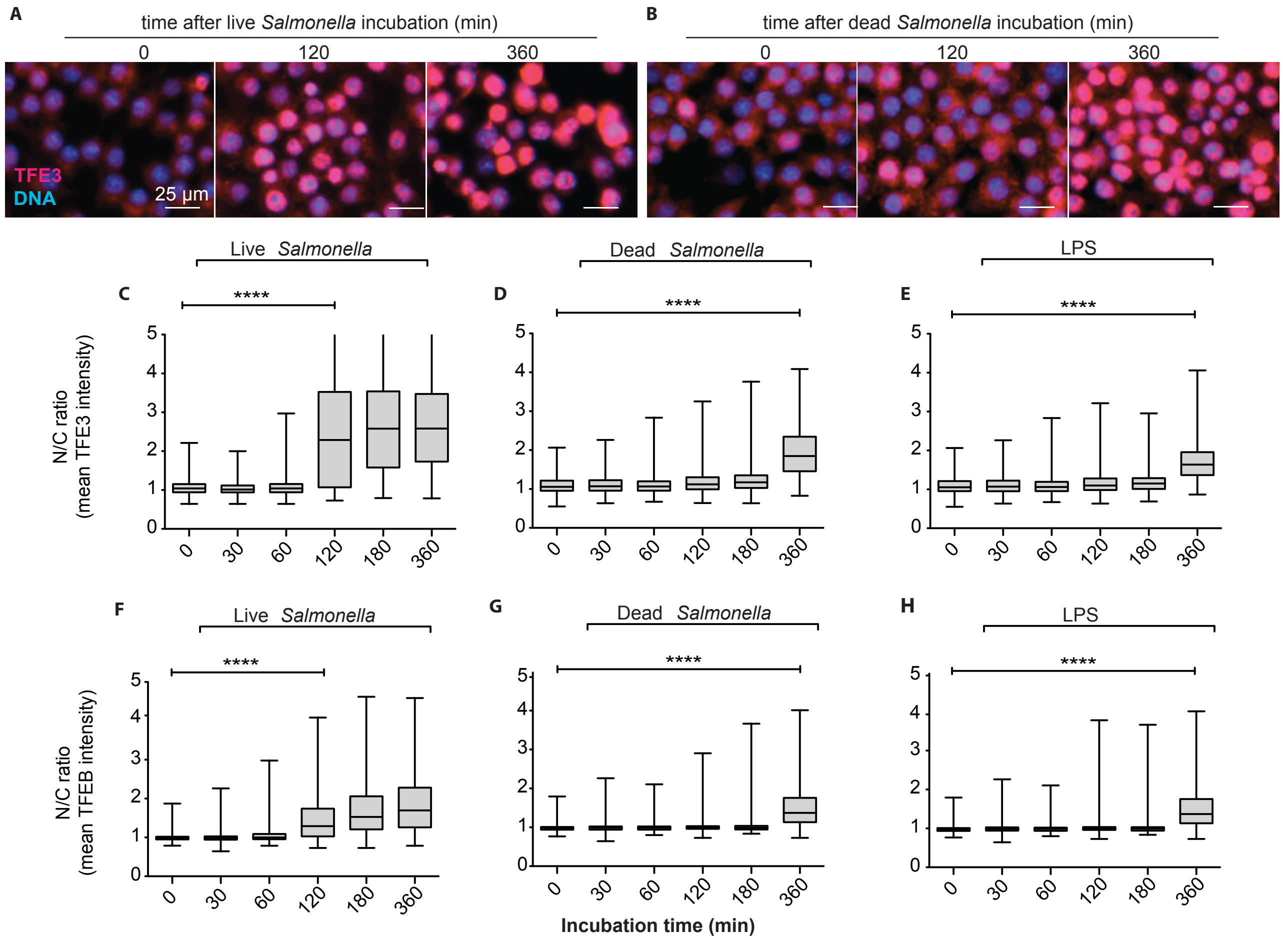

### Figure S2

Najibi et al - FIGURE S2

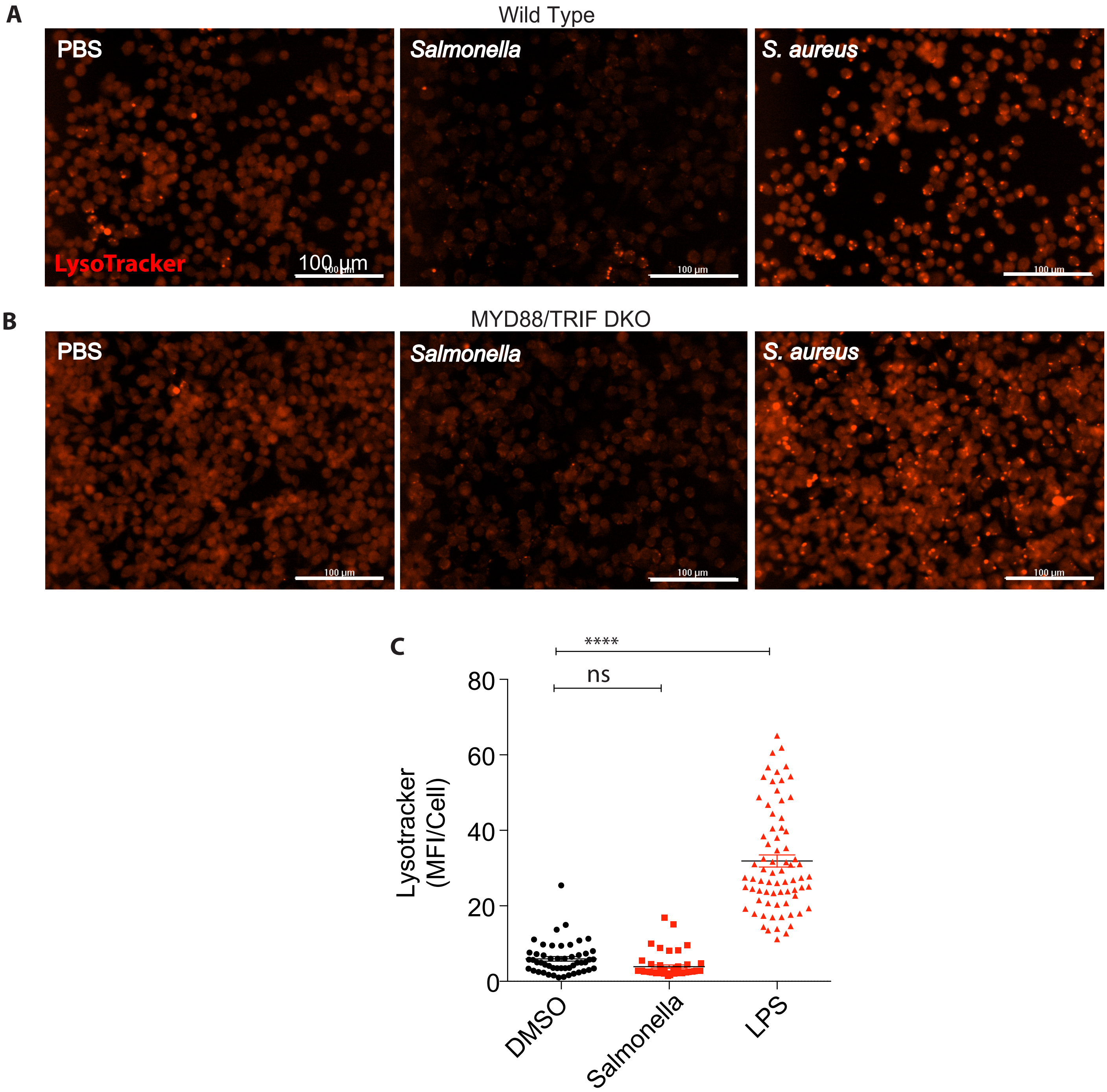

### Figure S3

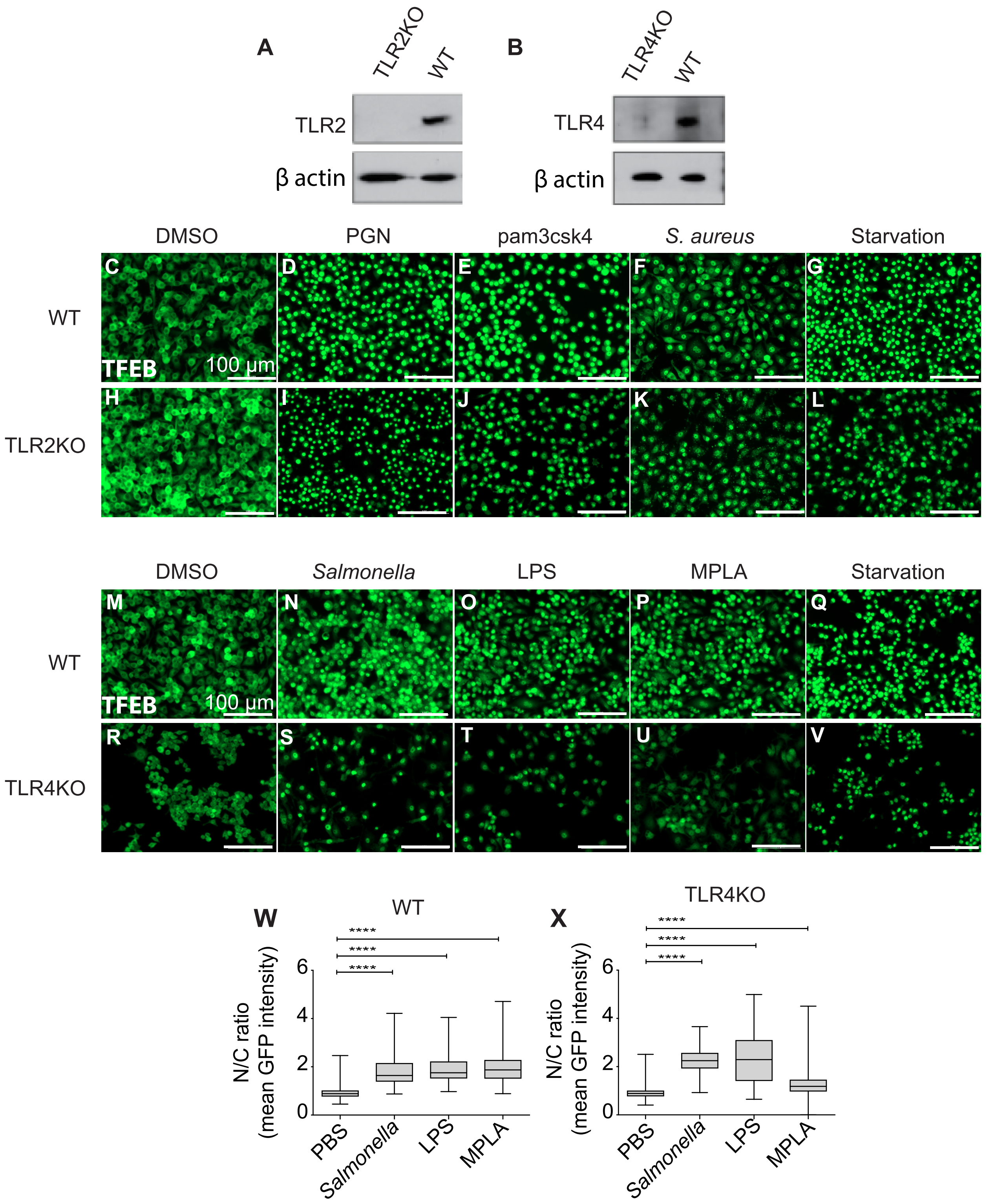

### Figure S4

Najibi et al - FIGURE S4

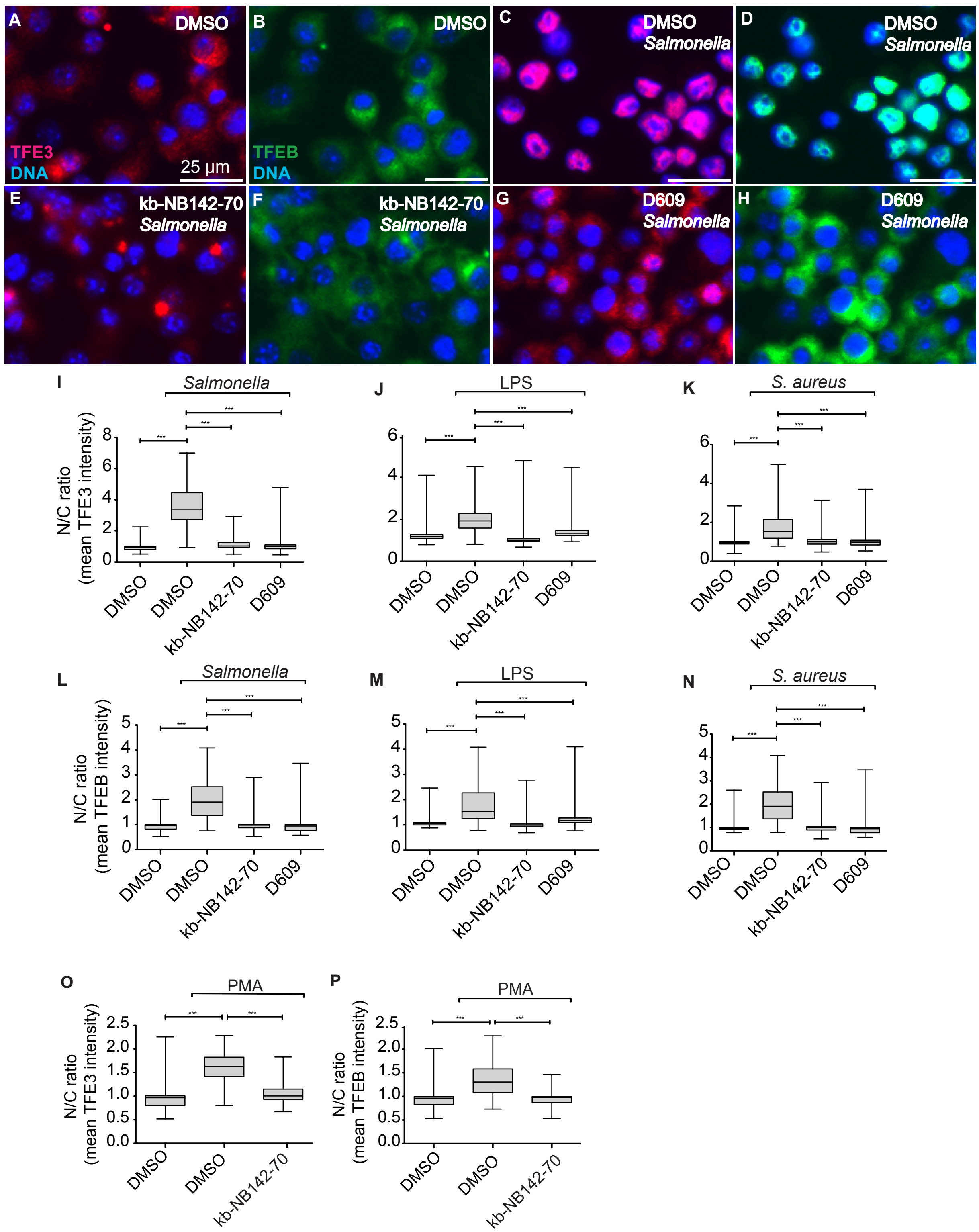

### Figure S5

Najibi et al - FIGURE S5

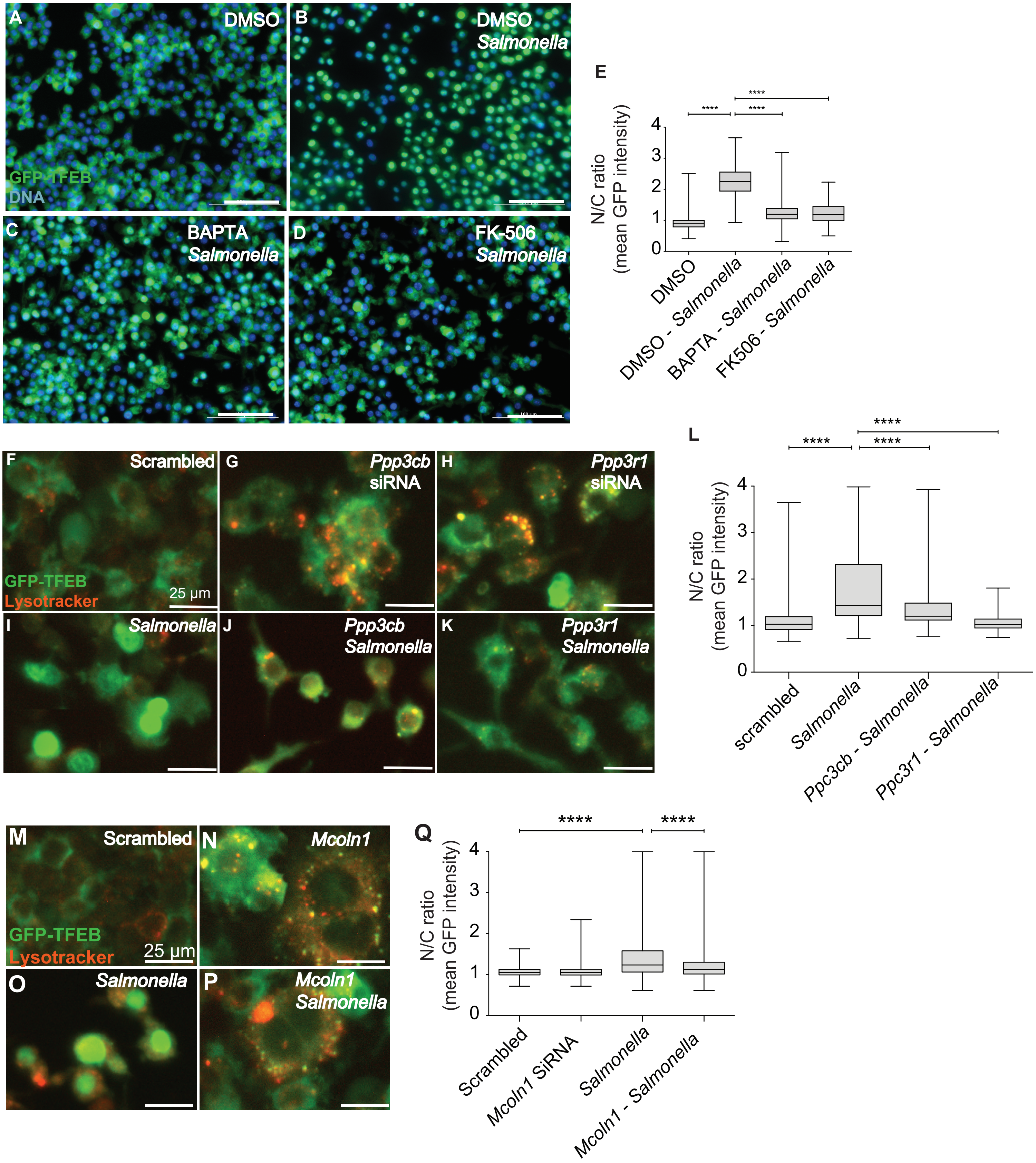

### Figure S6

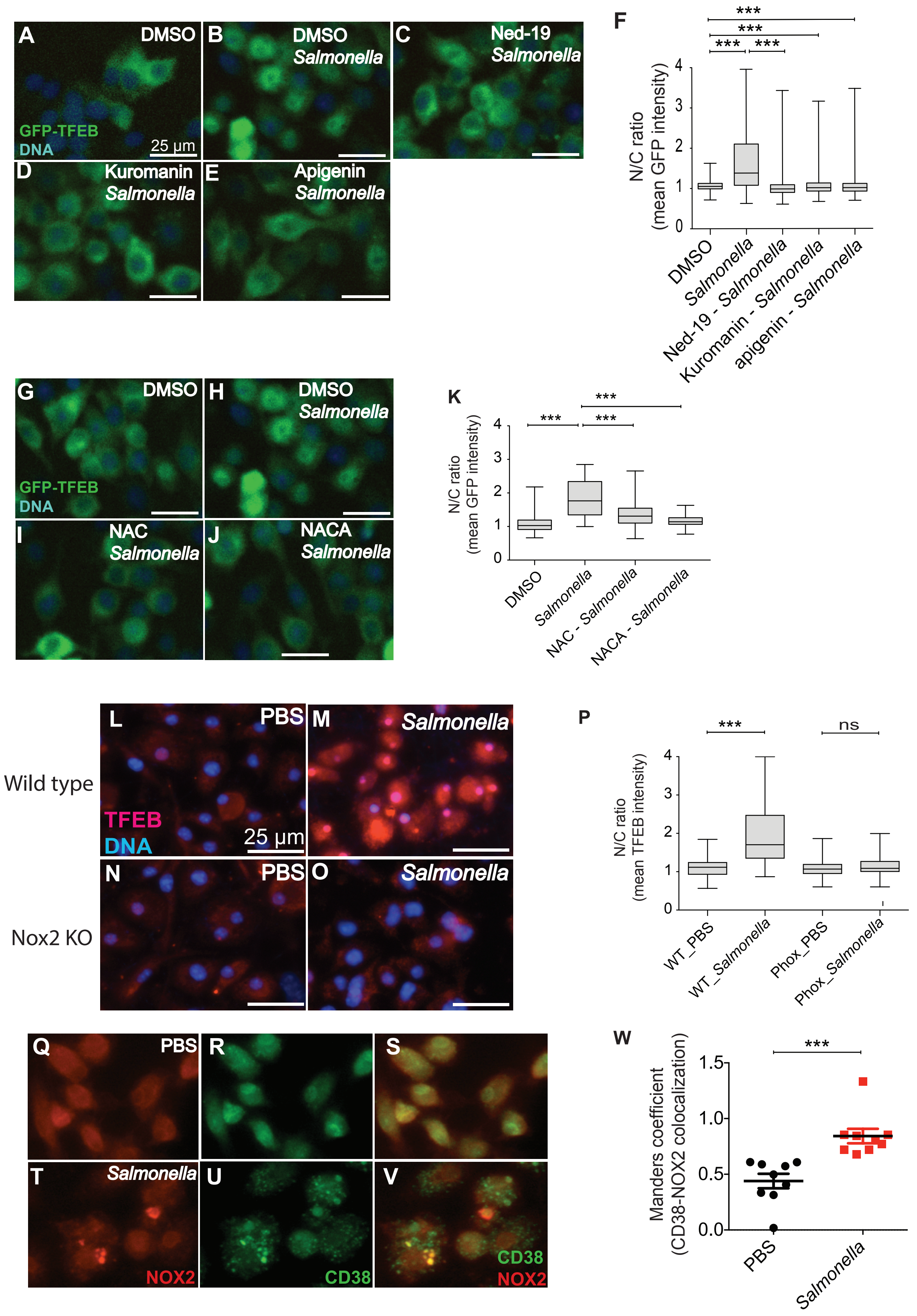

### Figure S7

Najibi et al - FIGURE S7

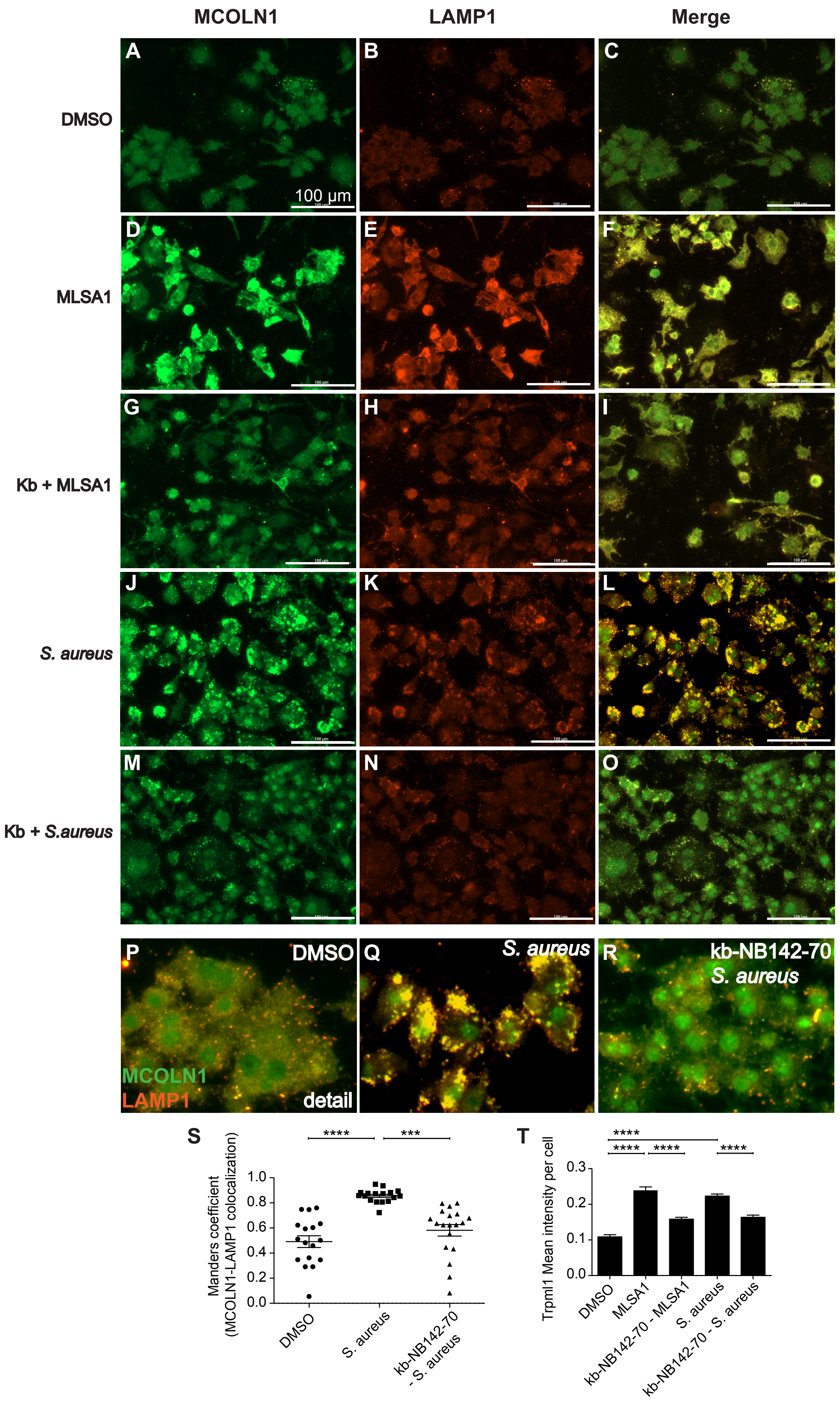
